# Initiation of primary T cell—B cell interactions and early antibody responses in an organized microphysiological model of the human lymph node

**DOI:** 10.1101/2025.01.12.632545

**Authors:** Djuro Raskovic, Jonathan M. Zatorski, Abhinav Arneja, Saweetha Kiridena, Tochukwu Ozulumba, Jennifer H. Hammel, Parastoo Anbaei, Jennifer E. Ortiz-Cárdenas, Thomas J. Braciale, Jennifer M. Munson, Chance John Luckey, Rebecca R. Pompano

## Abstract

In vitro microphysiological systems (MPS) are needed to replicate events in the lymph node (LN) leading to humoral immunity against new immune threats, but current lymphoid MPS focus largely on recall responses from memory lymphocytes. Here, an LN MPS was developed from primary, naïve human lymphocytes in microfluidic 3D culture to model interactions and antibody production at the LN T cell—B cell border. Naïve CD4+ T cells exhibited CCL21-dependent chemotaxis, chemokinesis, and activation in the MPS, and were skewed to a T follicular helper (pre-Tfh) phenotype. IgM secretion was induced in co-culture with activated B cells in the presence of a superantigen, staphylococcal enterotoxin B (SEB); micropatterning confirmed that the interaction required physical proximity. SEB-dependence of IgM secretion was greatest at a 1:5 T:B ratio, while seeding more pre-Tfh cells accelerated plasmablast differentiation and clustering. On-chip co-cultures at a 1:5 T:B ratio developed large lymphoid clusters containing CD38+ plasmablasts and CD138+ plasma cells after 15 days, with response varying between donors. Significant plasmablast induction in T-B co-cultures did not require the pre-Tfh phenotype, but pre-Tfh cells were required for inducing IgM secretion. We envision that this LN MPS will enable predictions and mechanistic analyses of human humoral immunity in vitro.

## INTRODUCTION

The lymph node (LN) is a small, intricately organized organ where antibody responses initiate after infection, vaccination, and the start of autoimmunity. Much of the action occurs in the border zones between the central T cell region and the B cell follicles (**Figure 1a**). Here, in response to antigen challenge and inflammatory signals, naïve T and B cells undergo a multi-step cascade of events and interactions in order to generate antibodies for the first time. Predicting these interactions in human lymph nodes is of great interest, with potential applications of in vitro models including drug and vaccine screening, immunotoxicology, and mechanistic models of human disease.^1^ However, prediction has been difficult due to limited access to culturable human lymphoid tissue and a lack of 3D cultured models that start from naïve T cells and B cells, even when using model antigens.

**Figure 1.**
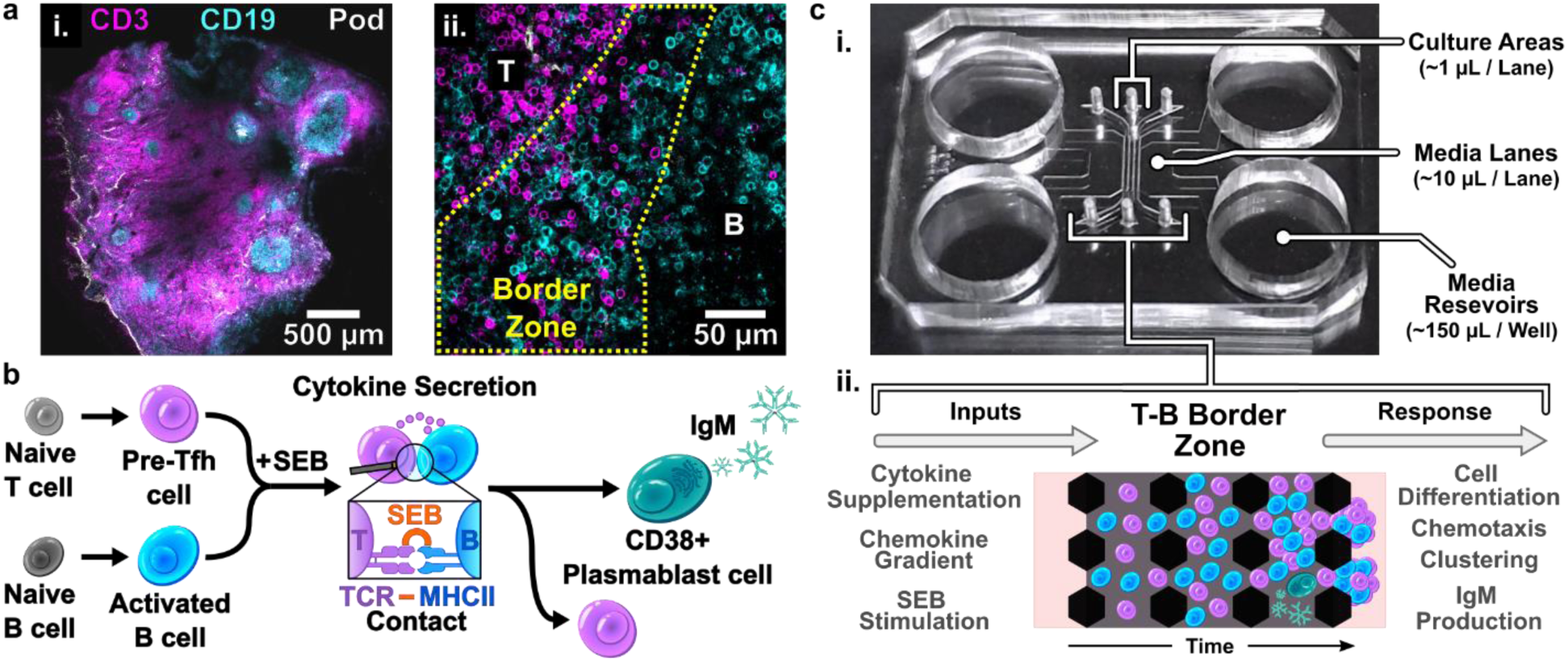
Approach for modeling early humoral responses to foreign protein using naïve T and B cells within the LN-MPS. (a) Immunofluorescence image of a human tonsil slice (i) with magnified view of the T-B border (ii). FITC-anti-Podoplanin, white; AF647-anti-CD3, magenta; PE-anti-CD19, cyan. (b) Illustration of biological events being recapitulated in the LN-MPS for modeling humoral immunity. Naïve T and B cells receive stimulatory signals that result in cell differentiation towards a pre-Tfh and activated B phenotype, respectively. Pre-Tfh cells provide help to activated B cells in a TCR-MHCII-dependent manner which is initiated upon exposure to cognate antigen or superantigen, such as staphylococcal enterotoxin B (SEB). Activated B cells mature into antibody-secreting plasmablasts. (c) Image of the LN-MPS chip with highlighted regions of interest (i) and illustration of model response to applied extrinsic factors (ii).

Reproducing T cell-mediated antibody production in vitro requires replicating a multi-step cascade of activation, differentiation, and physical contacts that occur early after an immune challenge. In the specialized LN microenvironment, stromal-derived signals such as Activin-A help drive naïve CD4+ T cells into T follicular helper cells (Tfh) upon T cell receptor (TCR)-ligation (**Figure 1b**).^2,3^ These T cells upregulate the chemokine receptor CXCR5, which enables them to migrate from the central T cell zone towards the border region near B cell follicles.^4^ Referred to as pre-Tfh cells, these early-activated CD4 T cells express Tfh phenotype markers (CXCR5, PD-1, and the transcription factor BCL-6) and can provide help to B cells in the border zone.^5,6^ Meanwhile, naïve B cells activated by inflammatory molecules upregulate expression of major histocompatibility complex class II (MHC-II),^7^ and also move to the B cell follicle border. There, interactions with cognate pre-Tfh cells drive differentiation of activated B cells into plasmablasts that secrete IgM, the first class of antibody secreted in a humoral response.^8,9^ This dance of contact-dependent, multistep interactions is thought to be critical for naïve lymphocytes to generate an effective initial humoral response to most novel, T cell-dependent antigens.

Much of this knowledge of basic LN immunology was derived from animal models, and with current technology it is difficult to assess which details hold true in human tissue. Mouse and human immune signaling, though broadly conserved, differ in numerous ways, including the response to TLR7/9/11 ligation,^10,11^ expression of lymphocyte-attractive chemokines,^12,13^ and mechanistic roles of some cytokines, such as IL-12.^14,15^ Meanwhile, clinical analysis of human immune function usually relies on direct analysis or 2D culture of blood draws.^16,17^ To better replicate tissue-level events, 3D culture systems have modeled interactions of lymphocytes with extracellular matrix, chemotaxis and organization,^18–22^ and pair-wise interactions such as dendritic and T cell engagement.^23–25^ However, models of multi-step tissue level events are rare.^1,26^ For higher-order processes, ex vivo slice cultures of lymphoid tissue and organoid models sourced from tonsil or blood have elegantly replicated B cell differentiation and antibody production in response to vaccination or infection.^27–32^ These systems are an exciting path forward for studies that do not require manipulation of the tissue microenvironment or imposed spatial organization, which are difficult to achieve in intact tissue or even an organoid. Integration of immune organoids with microfluidic perfusion and gradient culture is promising but in early stages.^29,33,34^

Lymphoid microphysiological systems (MPS) offer a promising alternative for additional environmental control. By incorporating a carefully chosen subset of organizational and functional features of in vivo tissue, MPS may predict specific tissue-level mechanistic events and responses to drugs.^35–38^ Development of immunocompetent MPS started over 10 years ago and has increased in the last few years, but only a handful of MPS specifically focus on the LN.^1,39–41^ Exciting progress has been made in modeling T cell-dendritic cell interactions,^42^ T cell-stromal cell interactions,^43,44^ vaccine responses and immunosenescence,^45,46^ and inflammatory and vaccine induced B cell clustering, memory B cell maturation, and antibody production.^34,47–49^ Engineered systems that permit imaging, including organoids^27,50^ and mixed 3D cultures in microchannels,^47^ have shown spontaneous self-organization of B cells and other cells into clusters.^29,47^ While powerful, these do not provide *a priori* control over the cellular organization or chemokine distribution in the culture, for example to test the role of spatial organization. While methods for patterning lymphocytes have been reported,^51–53^ including an elegant study with murine lymphocytes,^48^ a lymphoid MPS that supports patterning of human immune function has not yet been demonstrated.

Furthermore, in vitro lymphoid models of antigen-mediated immunity have focused primarily on recall responses, meaning responses to antigens encountered previously by the cell donor. Memory B cells respond readily to antigen, with less dependence on Tfh for support than naïve B cells.^54^ Prior exposure results in a high frequency of memory T cells and B cells that recognize that antigen, making it feasible for even a microscale MPS to generate a response to challenges with influenza, SARS-CoV-2, or other commonly encountered pathogens.^27,29,34^ In contrast, the precursor frequency of T cells specific to most novel antigens is extremely low (1 in 10^5^ to 10^7^).^55–57^ As a result, 3D cultured in vitro models have struggled to replicate the early activation and differentiation of naïve T cells and B cells and the subsequent interactions that result in antibody production. Besides antibody production from naïve B cells, even in vitro differentiation of naïve T cells into the required pre-Tfh state has never been specifically demonstrated in an MPS.

To address these gaps, here we describe a multi-lane microfluidic chip to model early T—B cell interactions at the border zone, starting with naïve cells and leading to antibody production (**Figure 1c**). The selected MPS format enabled either mixed co-cultures or well-controlled patterning of spatial organization. We tested the ability of the lymph node MPS (LN-MPS) to support cell activation and cell motility, differentiation into pre-Tfh cells, and multicellular T—B interactions. To sidestep the issue of low antigen specific precursor frequencies amongst naïve cells, we used the model antigen staphylococcal enterotoxin B (SEB), a bacterial superantigen. SEB has been used in traditional immunological cultures, and unlike standard antigen, reacts with most human leukocyte antigen proteins by crosslinking the TCR and B cell MHC-II, generating a polyclonal response.^58^ Consequently, most though not all donors should be capable of responding to SEB stimulation. Using this strategy, we optimized the LN-MPS to mimic T cell help for B cells, and tested cluster formation, plasmablast differentiation, and antibody production starting from naïve cells.

## RESULTS

### Design of the LN-MPS to provide options for mimicking lymphoid tissue spatial organization and function

When designing a LN-MPS to reproduce the activation, differentiation, and engagement of naïve T cells and B cells at the follicle border, we identified the following major design goals in terms of biological functions and analytical capabilities. Key biological functions included (a) naïve T cell activation and differentiation into cytokine-secreting pre-Tfh cells, (b) naïve B cell activation, (c) activated B cell differentiation into antibody-secreting plasmablasts, and (d) antibody production requiring T cell—B cell interactions. Required analytical capabilities included (a) control of lymphocyte organization in a 3D microenvironment, to mimic the overlap of T cell and B cell regions at the edge of the B cell follicle (**Figure 1a**); (b) spatial control over microenvironmental stimulation, such as to apply chemokine gradients; (c) spatially resolved analysis of cell motility and cell-cell interactions by imaging; (d) compatibility with immunofluorescence microscopy, sampling of cell secretions, and cell recovery for downstream analysis; and (e) ease of use by biomedical scientists.

To best model the naïve, human lymphocyte population in the MPS, primary lymphocytes were isolated from human blood TRIMA collars, which provide enriched leukocyte populations from patients following platelet donation. The use of TRIMA collars provided easy access to large quantities of autologous lymphocytes and circumvented any phenotypic variation associated with iPSCs or cell lines. To ensure that the chip did not contain effector and memory cells, we isolated naïve CD4+ T and naïve B lymphocytes using negative immunomagnetic selection for use on the chip. Although it is common to culture lymphocytes in media supplemented with fetal calf serum, here we chose serum free media to provide greater control over early activation states.^59^

We designed the LN-MPS to mimic the 3D tissue environment by resuspending the lymphocytes in a thermally-setting collagen-fibrinogen (Col/Fib) hydrogel.^60,61^ Three parallel gel lanes were implemented to provide flexibility for experimental design (**Figure 1c**). The well-established microfluidic format consists of parallel arrays of posts that provide surface-tension barriers for compartmentalized lane filling by simple pipetting.^62,63^ This planar design has been implemented routinely in organ-on-chip applications, is robust to use by non-experts, and is compatible with high-throughput manufacturing if needed in the future.^43,62,64^ The design supports cell loading in one, two, or all three lanes, either mixed or in separate populations, as well as patterning of distinct culture regions and application of chemokine gradients. As the three-gel-lane design is not yet available in an off-the-shelf format, we fabricated the device in PDMS with a glass bottom, which allowed for gas exchange and microscopic imaging, respectively.^65^ For simplicity and improved throughput, cells were cultured in static conditions throughout this work.

### The LN-MPS supported culture and activation of naïve T and B cells, and control over cell motility

As a first step, we tested the ability of the LN-MPS to support naïve T cell viability, activation, response to chemokines, and response to TCR ligation. Purified naïve CD4+ T cells were suspended in Col/Fib and loaded into the center lane of the chip, with the surrounding lanes filled with empty Col/Fib (**Figure 2a-b**). In preliminary experiments, larger dimensions of the gel lanes (3 x 800-μm-wide x 104-mm-long) caused limited gas exchange and poor viability of T cells (data not shown). A miniaturized configuration composed of 3 x 300-µm-wide channels with four reservoirs open for gas exchange (Figure S1) resulted in on-chip viability of naïve CD4+ T cells within 80% of that of 2D plated controls after four days across multiple donors (**Figure 2c**). IL-7 was supplemented in the media as it improved viability of naïve T cells in 2D off-chip experiments (Figure S2). A few T cells migrated spontaneously into the adjacent lanes, confirming motility in the hydrogel (**Figure 2b**). Chemotactic behavior was confirmed by adding CCL21 to the left media lane; within an hour, T cells migrated out of their initial lane toward the gradient (**Figure 2d-f**). Mean cell velocity was measured within expected ranges and showed expected chemokinetic responses to CCL21:^66^ ∼5 and ∼10 μm/min in the absence and presence of chemokine, respectively (**Figure 2g-h**). Responsiveness of naïve CD4+ T cells to ligation of the T cell receptor (TCR) complex was confirmed by injecting α-CD3/CD28 into both media lanes, resulting in IFN-γ secretion into the media over a period of 5 days (**Figure 2i**), visible enlargement of cells, and upregulation of the early lymphocyte activation marker CD69 at 48 hours (**Figure 2j-l**).

**Figure 2.**
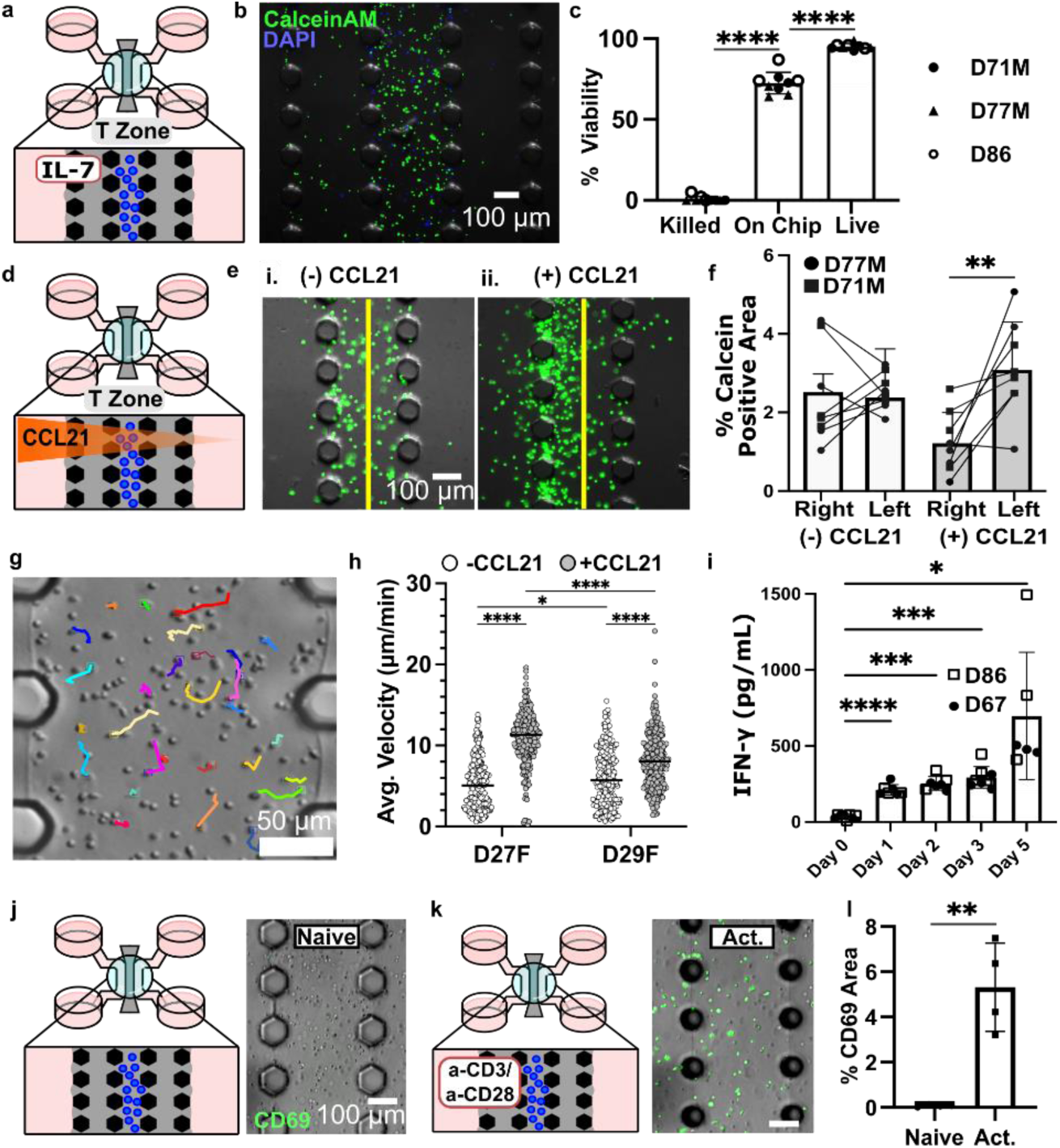
Naïve CD4+ T cells were viable, responded to an applied CCL21 chemokine gradient, and reacted to TCR-ligation within the LN-MPS. (a) Schematic of naïve T cells cultured on-chip with IL-7. (b) Representative image and (c) quantification of T cell viability after 4 days, labeled with Calcein AM, green/live, and DAPI, blue/dead, for 3 donors. One-way ANOVA Tukeys multiple comparisons test, **** p<0.0001 (d) Schematic of an applied CCL21 gradient established on chip, with CCL21 added to the left-hand media lane. (e) Representative images taken after 1 hr. without (i) or with (ii) a CCL21 gradient on chip and (f) quantification of naïve CD4+ T cell migration in response to presence or absence of CCL21. Cells were stained with Calcein AM, green. (g) Representative overlay of CD4+ T cell motility tracks with CCL21. (h) Quantification of mean cell velocity 30 min. after addition or omission of a CCL21 gradient, cells left unlabeled. Two-way ANOVA Tukeys multiple comparisons test, * p<0.05, **** p<0.0001. (i) Quantification of IFN-γ secretion by activated or naïve CD4+ T cells on-chip measured by ELISA of supernatants collected on days 0 (naïve) and 5. Mixed effect analysis, Dunnett’s multiple comparisons test, * p=0.03, *** p=0.0001, **** p<0.0001. Schematic of naïve T cells cultured without (j) and with (k) a-CD3/CD28 with images of CD69+ signal (FITC-anti-CD69, green) for (j) naïve and (k) TCR-stimulated (Act.) T cells on-chip, and quantification (l) of CD69 signal after 48 hours. Each dot represents a chip from D71M. Statistics determined using an unpaired T test, ** p<0.005.

In a separate experiment, we verified that the LN-MPS supported B cell receptor (BCR)-mediated B cell activation in monoculture. We used a skewing cocktail composed of α-IgG/IgM for BCR ligation, R848, a TLR agonist, and CD40L, which mimics T cell help. As expected, the activated B cells became visibly larger and increased expression of CD69 and CXCR5 compared to naïve controls (Figure S3). Thus, the LN-MPS supported naïve T and B cell stimulation, and analysis of critical primary CD4+ T cell functions including chemotactic behavior.

### Naïve T cells differentiated into early T follicular helper cells on chip

Pre-Tfh cells are critical for support of B cell maturation and T cell-dependent antibody production. These cells are characterized by upregulation of the chemokine receptor CXCR5, regulatory protein PD-1, and the master transcription factor BCL-6 compared to naïve T cells, as well as secretion of IL-21.^4^ For human cells, TCR engagement in the presence of cytokines interleukin-12 (IL-12) and activin A are capable of driving naïve T cells to pre-Tfhs.^3^ Here we sought to implement pre-Tfh differentiation from naïve T cells in an MPS for the first time.

To assess skewing to a pre-Tfh phenotype, naïve CD4+ T cells were loaded into the chip and cultured for 5 days with a “skewing cocktail” composed of IL-7, α-CD3/CD28, IL-12, and activin-A, added to the media lanes (**Figure 3a**). Cells in the skewed condition formed small clusters in the hydrogel, which were CXCR5+ by immunofluorescence staining; little CXCR5 signal was observed outside of the clusters or in the unstimulated controls (**Figure 3b**). T cells treated with the skewing cocktail secreted significantly more IL-21 than naive controls (**Figure 3c**). To further characterize the T cell state, cells were recovered from the chip after gel digestion by collagenase and analyzed by flow cytometry (**Figure 3d**), which showed induction of CXCR5+, PD-1+, and BCL6+ across three donors. Gating strategies are provided in Figure S4. Thus, naïve CD4+ T cells were successfully skewed to a pre-Tfh state in the LN-MPS. In later experiments, we found that inclusion of IL-7 was not required for promoting T cell viability during pre-Tfh differentiation (data not shown), and omitted this cytokine from the cocktail.

**Figure 3.**
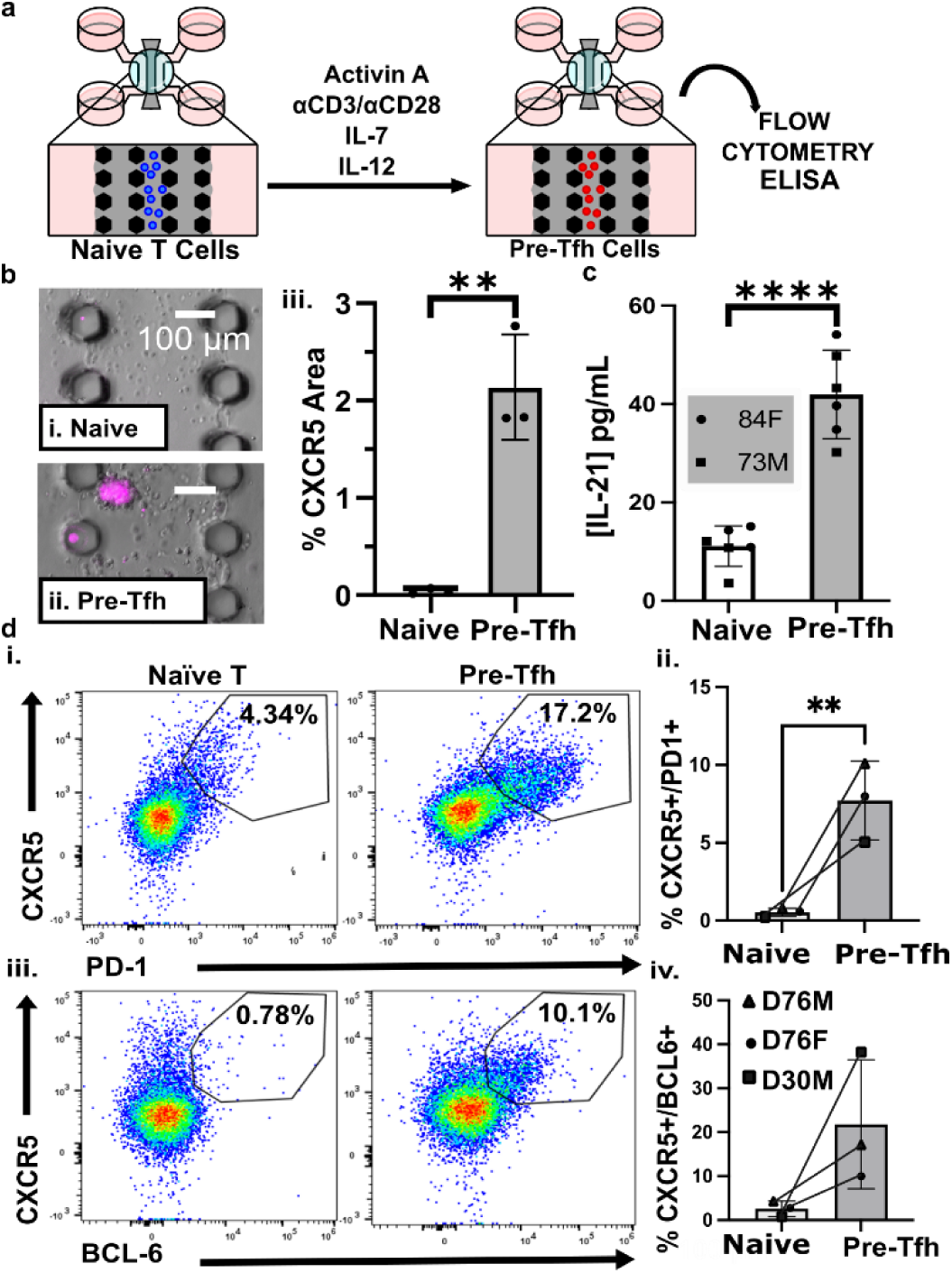
The LN-MPS replicated human pre-Tfh development. (a) Schematic of naïve T cells cultured on-chip with a-CD3/CD28 and a cytokine cocktail (IL-7, IL-12, and Activin A) to induce differentiation into a pre-Tfh phenotype. (b) Overlays of immunofluorescence (AF647-anti-CXCR5, magenta) and brightfield images after 3 days culture for one representative donor, 71M, (i) with or (ii) without pre-Tfh-skewing cocktail. (iii) Quantification of percent CXCR5 area across each image. Each dot shows the mean from one chip. Unpaired t test, ** p<0.005. (c) IL-21 concentration from chip supernatants, quantified by ELISA. Unpaired t test, **** p<0.001. (d) Flow cytometry characterization of pre-Tfh markers from cells recovered from chips. (i) Flow plots with (ii) quantification of the percent CD4+ T cells expressing CXCR5 and PD-1. (iii) Flow plots with (iv) quantification of the percent CD4+ T cells expressing CXCR5 and BCL6. Each dot is pooled from 20 chips/donor. Ratio paired T test, ns p>0.05, * p<0.02. Bar graphs show mean +/- standard deviation.

### SEB induced engagement between pre-Tfh and activated B cells resulted in antibody secretion that required close proximity

Next, we tested the extent to which the LN-MPS could predict B cell antibody secretion in response to the model antigen staphylococcal enterotoxin B (SEB), which allows polyclonal interactions between the two cell types. To precisely control the numbers of live cells loaded into the LN-MPS, naïve T and B cells were skewed individually in 2D into their pre-Tfh and activated B states prior to loading with or without SEB (**Figure 4a**). Drawing inspiration from Wagar et. al,^27^ an insulin, transferrin, and selenium (ITS) supplement was added in all conditions to support viability in the T-B co-cultures (Figure S5). Furthermore, we exploited the patterning capabilities of the LN-MPS to test the hypothesis that in the absence of any chemokine gradients or stromal cues, pre-Tfh and activated B cells would require close physical proximity to produce antibodies in vitro. Pre-Tfh and activated B cells were loaded either into individual outer lanes or combined in the central lane at a 1:1 ratio (**Figure 4b**). On day 6, IgM secretion was dependent on both the presence of SEB and close T-B cell proximity (**Figure 4c**). In contrast, on day 3, IgM secretion was not SEB-dependent, but was still dependent on seeding proximity. (**Figure 4d**, S6). These data showed that pre-Tfh cells assist B cells with IgM secretion in the LN MPS, but indicated that optimization was required to increase the dependence on SEB-mediated engagement.

**Figure 4.**
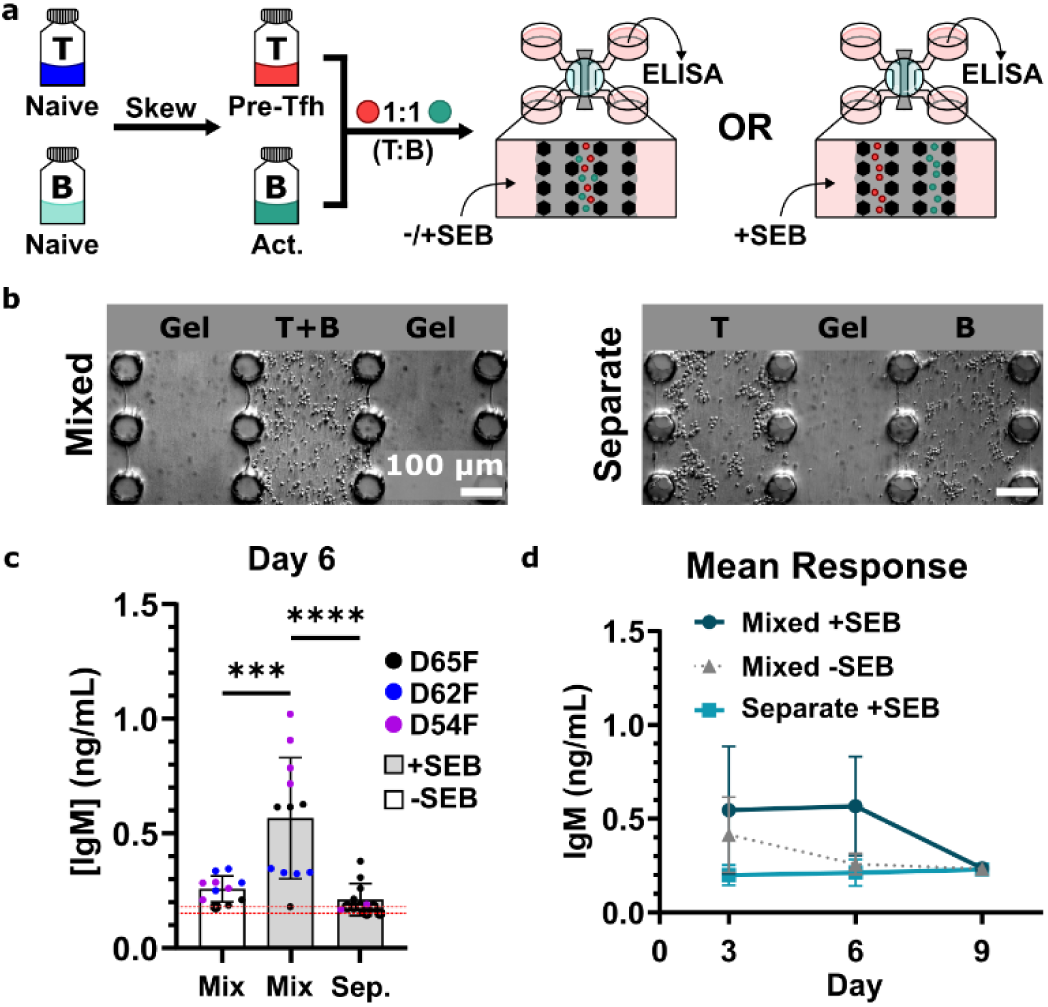
IgM secretion was dependent on the proximity of pre-Tfh and activated B cells when co-cultured in the LN-MPS. (a) Schematic of experimental setup for analyzing contact-dependent pre-Tfh induction of IgM secretion by activated B cells. Cells were skewed in 2D monoculture for 3 days prior to seeding in the LN-MPS. T and B cells were either seeded together in the central lane, with or without SEB added to the culture media, or in the outer lanes separated by a lane of empty col/fib hydrogel, with SEB added to the culture media. (b) Representative images from D65F showing spatially patterned cells within the LN-MPS on day 3. Cell distribution and density was similar on day 6 and 9, not shown. (c) Quantification of IgM secretion for day 6. Each dot shows a single chip replicate. Bars show mean and standard deviation across three donors. Dashed red line is the limit of detection. *** p=0.0002, **** p<0.00001, statistics determined using an ordinary one-way ANOVA with Sidak’s multiple comparisons test with single pooled variance. (d) Quantification of IgM secretion across the 9 day co-culture period. Each dot represents the mean response of three donors for a given condition, and error bars show the standard deviation across donors.

### Clustering, plasmablast formation, and IgM production in response to SEB depended on the cell seeding ratio

In vivo, the number of T cells in a follicle correlates linearly with the number of B cells,^67^ suggesting that the local ratio of T to B cells may be a critical parameter for immune function. Indeed, in classic 2D co-cultures, the ratio of human pre-Tfh cells to B cells significantly impacted measured outcomes,^68^ but it is not possible to test this prediction in vivo, and no 3D cultured system has been available to do so. We hypothesized that LN-MPS would provide a means to explore the impact of human T:B ratio on the magnitude of T cell-dependent humoral responses. Therefore, we investigated the effect of the pre-Tfh:B cell ratio on IgM secretion, plasmablast (CD38+) formation, and emergent self-organization after SEB stimulation. Once again, to precisely control the ratios of these cells in the LN-MPS, we pre-skewed naïve T to pre-Tfh and naïve B to activated B cells in separate 2D cultures (off-chip). Then, keeping total density constant, pre-Tfh and activated B cells were mixed at ratios of 1:1, 1:5, and 1:10, respectively, in gel precursor solution, loaded in all three lanes of the LN-MPS, and cultured with or without SEB (**Figure 5a**).

**Figure 5.**
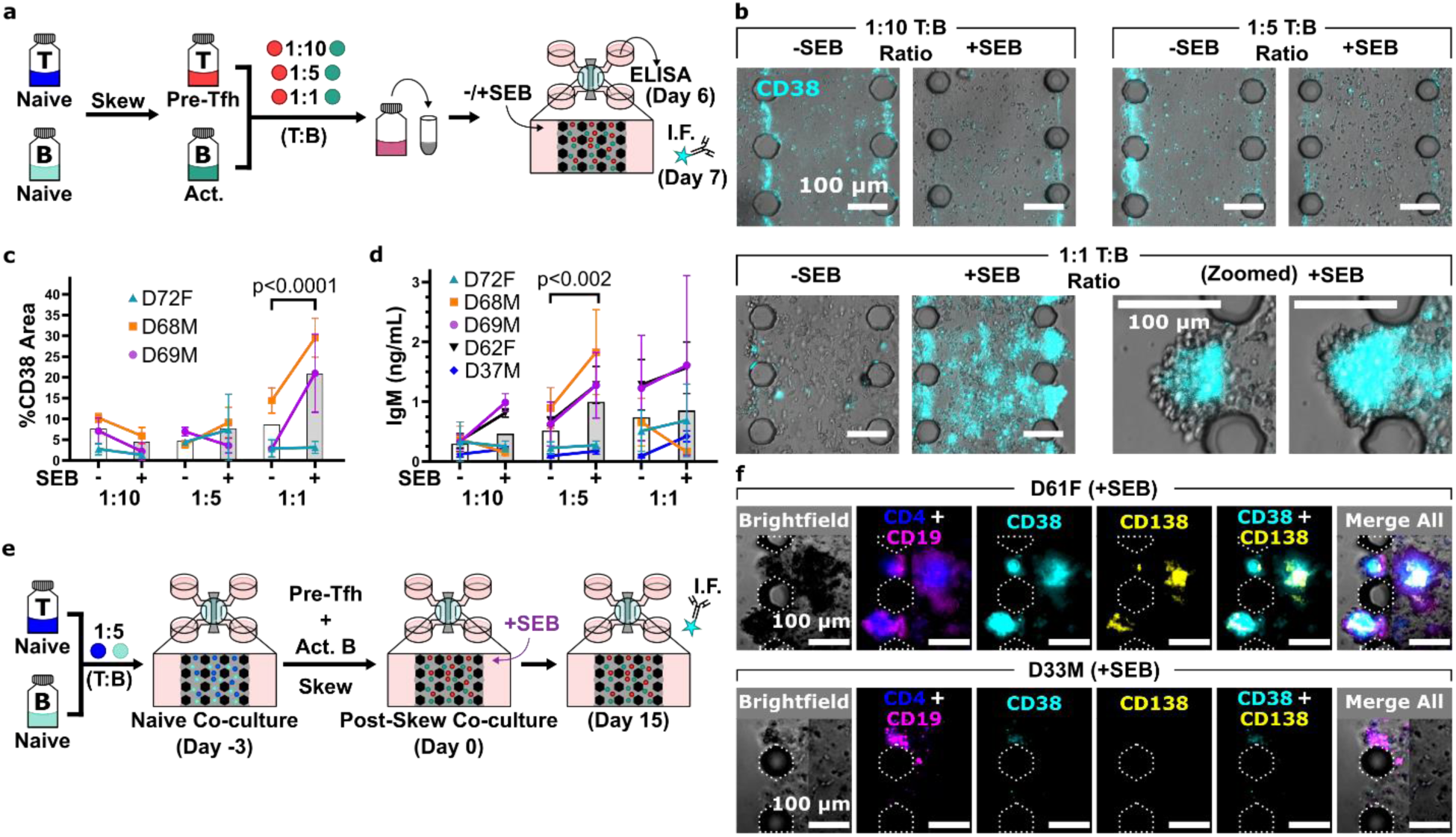
SEB-dependent IgM secretion and B cell maturation towards a CD38+ plasmablast-like and CD138+ plasma cell-like phenotype were decoupled and sensitive to initial cell seeding conditions in the LN-MPS. (a) Schematic of experimental setup for evaluating the impact of different T:B cell seeding ratios on IgM secretion and B cell maturation towards a CD38+ plasmablast-like state. Cells were skewed in 2D monoculture and then suspended in a col/fib hydrogel at a 1:10, 1:5, or 1:1 T:B ratio. Cell suspensions were placed in the LN-MPS and cultured with or without SEB for 7 days. IgM secretion was measured by ELISA on day 6, and immunofluorescent (I.F.) staining was performed on day 7. (b) Representative images from D69M showing CD38+ plasmablast-like cells, stained with AF546-anti-CD38, cyan. (c) Quantification of percent positive CD38 area across the whole image (3 donors, 2-6 chips/donor). Statistics determined using mixed-effects model with restricted maximal likelihood and matched values spread across rows. (d) Quantification of secreted IgM as measured by ELISA. Statistics determined using repeated measures two-way ANOVA with matched values spread across rows with Šidák’s multiple comparisons test and single pooled variance. For panels c and d, each dot is the mean signal for a given donor with error bars showing standard deviation of the signal for a given donor. Background bars represent the average response across all donors. (e) Schematic of experimental setup for evaluating the impact of skewing cells in 3D co-culture, and maintaining the co-culture for a longer duration of time. Naïve T and B cells were suspended in a col/fib hydrogel at a 1:5 ratio and placed in the LN-MPS with a combined T-B skewing cocktail and ITS added to the media lanes for 3 days. On day 3, skewing cocktail containing media was removed and chips were rinsed following addition of the media with SEB. (f) Representative images showing variable donor response with regards to B cell maturation when cells are skewed in 3D co-culture. Dotted outlines indicate posts. Cell stained with BV421-anti-CD4, blue; AF647-anti-CD19, magenta; AF546-anti-CD38, cyan, DyLight488-anti-CD138, yellow.

Unlike the single central lane of cells in **Figure 4**, the three lanes of cells at a 1:1 T-B ratio resulted in IgM production at day 6, apparently regardless of the presence of SEB, suggesting potential overstimulation of B cells in this condition (**Figure 5b-c**). At this ratio, large self-organized clusters spontaneously formed in response to the addition of SEB, and were present between posts and jutted out into the media lanes in an organoid like fashion. Fascinatingly, formation of CD38+ plasmablast-like cells was dependent on SEB at this density, and furthermore the CD38 signal was particularly concentrated within the cell clusters at the edge of the gel, rather than in the central gel regions.

Interestingly, the magnitudes of IgM secretion (**Figure 5d**) and plasmablast formation were decoupled. Whereas the greatest SEB-dependence of CD38+ signal was measured at a 1:1 T:B ratio, the greatest SEB-dependence of IgM production was measured at a 1:5 T:B ratio. Little response was observed at the 1:10 T:B ratio at this timepoint. These data were consistent with a catalytic function of pre-Tfh cells for B cells. We theorize that in a 3D microenvironment with a 1:10 T:B ratio, B cells rarely encounter a pre-Tfh cell and thus fail to mature into antibody-secreting plasmablasts. Meanwhile, 1:1 T:B ratios provide maximal support for rapid B cell maturation towards a CD38+ plasmablast-like state. However, the high cell density caused consistent cell-cell contact, resulting in overall enhanced secretion of IgM, but loss of SEB dependence. Meanwhile, on average a 1:5 T-B seeding ratio provided the optimal conditions for frequent, but controlled, instances of TCR-MHCII receptor engagement between T and B cells, thereby recovering SEB dependence for IgM secretion.

Variation in the magnitude of response to antigen challenge is expected in the human population, and similar variation was observed in the responses to SEB on the LN-MPS. Out of five donors tested, three responded to SEB with increased IgM secretion at the optimal 1:5 T:B ratio, while two donors were largely non-responsive with at most a small response to SEB at the highest T:B ratio (1:1). Exploration of this population variation in humoral immunity is an exciting future application of the LN-MPS. Since antibody secretion is the ultimate contributor for assessing the quality of protection provided by the humoral immune response, we opted to use this metric for determining optimal conditions for the LN-MPS, and proceeded with a 1:5 T-B seeding ratio for future experiments.

### Plasma cell formation was possible in LN-MPS held for longer duration

We hypothesized that 1:5 seeding ratios could potentially achieve high cell densities, similar cluster formation, and further B cell maturation in chips held for longer times. We assessed this in a separate experiment in which naïve T and B cells were skewed in situ in co-culture within the LN-MPS (1:5 ratio), rather than separately prior to loading, for reduced handling of the culture (**Figure 5e**). We maintained the LN-MPS for a total of 18 days: 3 days for skewing and 15 days with SEB. Cultures were visually monitored for formation of dense clusters at the gel-media interface, followed by immunofluorescence staining and imaging on day 15. Qualitatively, we observed that, on average, cell density increased within the central hydrogel areas for approximately 9 days after skewing, after which it began to plateau (Figure S7). Meanwhile, dense clusters formed at the gel-media interface around day 9, and began to plateau around day 12 post-skew. At day 15 post-skew, immunofluorescent staining showed dense CD38+ plasmablast-like and CD138+ plasma cell-like groups in some but not all donors (3 donors tested), similar to the variation observed in IgM secretion for earlier experiments (**Figure 5f**). As before, the CD38 and CD138 signal were most frequent at the gel-media interface. We conclude that the 1:5 T:B ratio supported the generation of plasmablast (CD38+) and plasma cell (CD138+) phenotypes in the LN-MPS, and that cultures could be maintained for at least 18 days on chip. Further testing is required to optimize the conditions and determine frequency in plasma cell response across the donor population.

### IL-21 and IgM secretion was dependent on both T cell phenotype and SEB in the LN-MPS, while plasmablast differentiation was dependent on SEB but not Tfh phenotype

Having determined an optimal cell seeding ratio, we tested the hypothesis that the pre-Tfh phenotype was required for optimal production of antibodies and differentiation of B cells in this system. In accordance with prior reports, we expected that relative to their naïve or activated (TCR ligation only) counterparts, pre-Tfh cells would act as enhanced inducers of plasmablast differentiation and antibody secretion due to their enhanced production of costimulatory molecules such as CD40L and enhanced production of IL-21.^6^ To assess this, naïve T cells were either maintained in their naïve state, activated using only αCD3/28, or skewed towards a pre-Tfh state with αCD3/28 supplemented with IL-12 and Activin A. Separately, naïve B cells were activated using R848 and αIgG/IgM, then loaded into the LN-MPS with the appropriate T cell population (1:5 T:B ratio). (**Figure 6a**).

**Figure 6.**
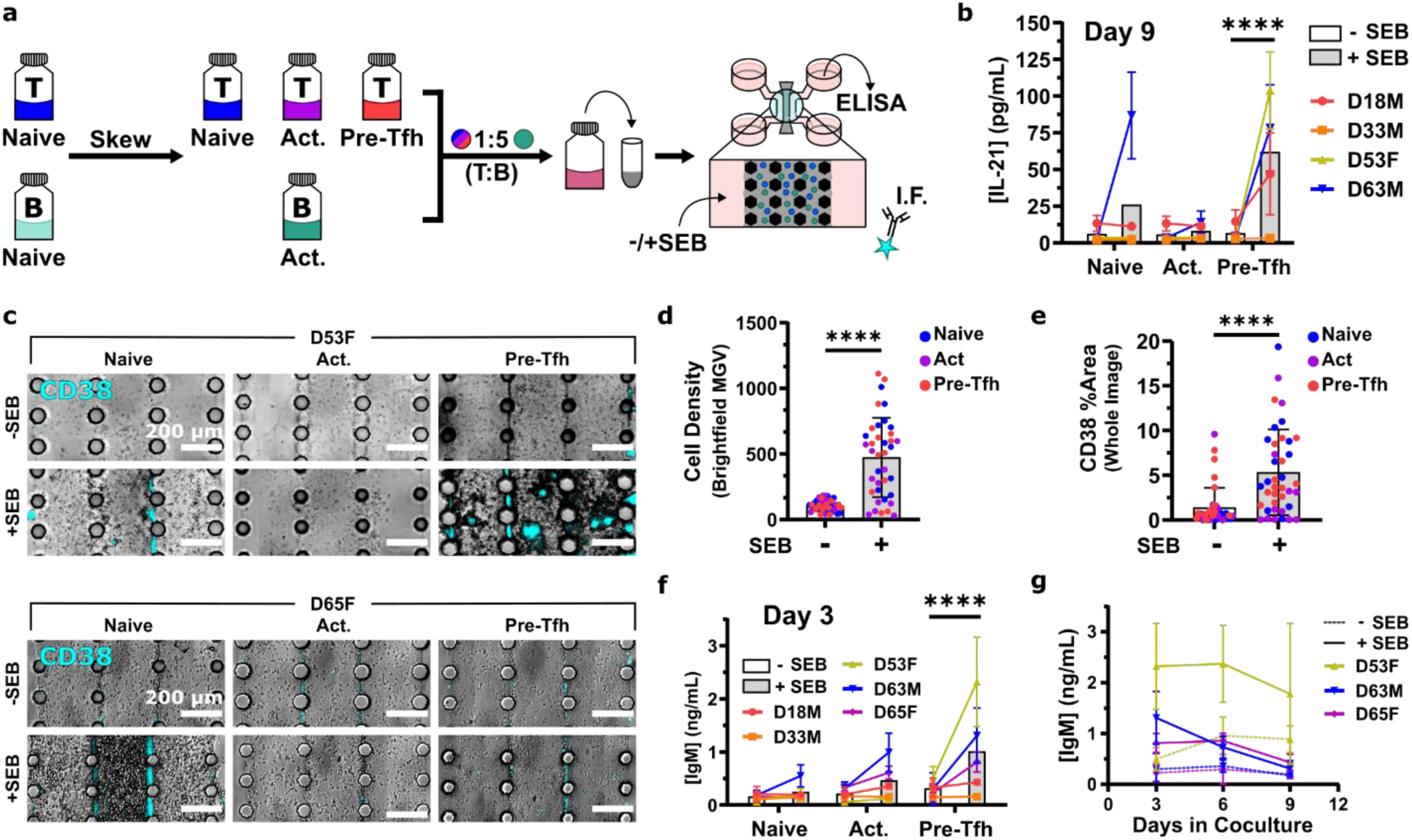
IL-21 and IgM secretion were dependent on the presence of pre-Tfh cells and SEB, whereas differentiation of CD38+ plasmablast-like cells was independent of pre-Tfh. (a) Schematic of experimental setup for evaluating impact of T cell phenotype on antibody secretion and B cell maturation within the LN-MPS. Cells were skewed in 2D monoculture for three days before suspension in col/fib hydrogel and seeding on-chip at a 1:5 T:B ratio. Media was added with or without SEB and the culture was maintained for 9 days. (b) Quantification of IL-21 secretion by ELISA on day 9 for each condition. (c) Representative images showing increased cell density at end-point analysis and variable degrees of CD38+ plasmablast-like cell clusters, AF546-aCD38, cyan, for each condition. (d) Quantification of increased cell density at end-point analysis in response to SEB using the absolute value difference between the MGV of the empty media lanes and cell-populated gel lanes. (e) Quantification of positive CD38% area in response to SEB. (f) Quantification of IgM secretion for each condition on day 3 with (g) accompanying timeline of IgM secretion for top three responding donors. Statistics for panels b and f determined using ordinary two-way ANOVA with Šidák’s multiple comparisons test and single pooled variance, *** p=0.0005, **** p<0.00001. Each dot is the mean response for a given donor measured from 3-4 individual chips, error bars show the standard deviation for a given donor, and background bars show the combined mean response. Statistics for panels d and e determined using an unpaired two-tailed t-test, **** p<0.00001. Each dot is the signal measured from an individual chip, error bars show the standard deviation, and background bars show the combined mean response (N=4 donors, 2-4 chips per donor per condition).

As expected, cocultures containing pre-Tfh cells secreted the greatest quantity of IL-21 even at day 9 (**Figure 6b**). TCR-ligation by αCD3/28 alone was not sufficient to induce detectable IL-21 secretion. Interestingly, IL-21 secretion was strongly SEB-dependent; cultures without SEB had little IL-21 in the media at day 9 (**Figure 6b**). Since SEB induces robust T-B cell interactions via crosslinking MHCII to TCR, these data are consistent with a known role of persistent cell-cell interactions between T and B cells being required for maintenance of pre-TFH phenotypes.^6^

As observed previously, both cell density and CD38 immunofluorescence increased significantly in response to SEB when all conditions were combined (**Figure 6c-e**). Interestingly, all CD4 T cell populations, not just the pre-Tfh, induced the increase in cell density and CD38 expression in the presence of SEB, though mostly not in its absence. These data suggest that the conditions used during off-chip B cell stimulation led to a hyperresponsive B cell population, one that was somewhat less dependent on Tfh-specific factors such as CD40L surface expression and IL-21 secretion, but still required T cell crosslinking via SEB for maximal effect. We speculate that this state may have resulted from the choice or concentration of TLR7/8 agonist, R848, during initial B cell activation protocol. An exciting future application of the LN-MPS is to analyze the sensitivity of the T-dependent B cell response to different TLR agonists and B cell activation conditions, with the goal of better mimicking different infections as they occur in a tissue-like setting.

In contrast to clustering and plasmablast differentiation, IgM secretion was strongly dependent on the state of the T cells, with the greatest IgM in co-cultures containing pre-Tfh cells and the lowest from naïve T cells (**Figure 6f-g**). IgM secretion was also dependent on the presence of SEB even at early timepoints, day 3, and was sustained up to day 6. On average, by day 9, IgM secretion was reduced to near that of the negative-control condition (no SEB) across all conditions (Figure S8b). As before, magnitude of IgM secretion and magnitude of CD38 signal were decoupled. It is possible that this was the result of performing immunofluorescent staining at a timepoint when antibody secretion was reduced, or it may indicate that CD38 expression is not directly correlated with antibody production. We conclude that the LN-MPS predicts the expected requirement of pre-Tfh cells for inducing IgM secretion in an SEB-dependent manner, whereas plasmablast formation was induced by both Tfh and non-Tfh CD4+ T cells in an SEB dependent manner.

## CONCLUSIONS

Here we present a primary human LN MPS to model T cell—B cell interactions at the lymph node T-B border. The system was designed for use by individuals with traditional benchtop expertise, requiring no microfluidic pumps or tubing, and is compatible with brightfield and fluorescence imaging. Using autologous naïve CD4+ T cells and B cells isolated from blood, the model reproduced expected functions of T cells and B cells. These included chemokinesis and chemotaxis of naïve T cells towards a CCL21 chemokine gradient, skewing of naïve T cells towards a pre-Tfh state, and activation of naïve B cells. Furthermore, the LN-MPS successfully replicated IgM secretion by B cells in a TCR-MHCII-dependent manner and required the presence of pre-Tfh cells. Inducing IgM secretion in response to SEB required close spatial proximity of pre-Tfh cells to activated B cells. The LN-MPS provided a platform to test the role of T:B cell ratio in a human 3D culture environment, where the 1:10 T:B co-culture ratio commonly used in 2D experiments was insufficient for plasmablast generation or antibody secretion. Instead, a 1:5 T:B ratio was required for TCR-MHCII-dependent IgM secretion, and in multi-week cultures it was sufficient to generate organoid-like clusters containing plasmablast-like (CD38+) and plasma cells (CD138+), with donor-to-donor variation in response.

The current MPS has several limitations that present opportunities for further advancement. The LN-MPS successfully modeled T cell chemotaxis towards CCL21, but due to the small length scale and typical rates of protein diffusion, we estimate that gradients remain intact for less than 3 hours prior to equilibration. Long-term gradient maintenance in microfluidic systems is traditionally accomplished by maintaining constant fluid flow, which could be introduced in future work. Here, fluid flow was deliberately omitted to enable easy scaling of the system to dozens or hundreds of chips simultaneously, but future incorporation of high-throughput pumping systems or gravity driven flow also would enable study of the effects of interstitial fluid flow. Finally, while the current system showed IgM production, we did not detect IgG in the supernatant. We speculate that additional biological cues may be required to induce class switching when starting with naïve B cells. Stromal cells and antigen presenting cells were omitted here for simplicity, but future studies including these cells are warranted to determine the cellular and molecular requirements for class-switching in this system. For example, stromal cells provide both a physical network for T and B cells interactions and immunoactive signals such as Activin A, IL-6, and chemokines.^69,70^ Their inclusion in future iterations of the LN-MPS may enhance the rate, magnitude, and duration of the antibody responses.

With its current focus on predicting IgM responses from naïve lymphocytes as well as with future expansions of the system, we envision the LN-MPS will enable many future investigations into human immunity. Differences in immune function and regulation between cells drawn from varying donor age groups, sexes, body masses, and other biological factors can easily be evaluated in future studies using this system. The system is also well suited to test the impact of immunostimulatory and suppressant drugs on T cell—B cell interactions and IgM production, both for drug testing and immunotoxicology. Furthermore, the LN-MPS allows for experiments to probe the role of organization at a tissue-level scale. The multi-lane platform provides control over regional cell organization, seeding ratios, densities, and establishment of short-term gradients of soluble factors, coupled with the ability to image on chip or collect samples for down-stream analysis. In the future, the system may also be coupled with other organ-on-chip systems to study the impact of upstream organs on T—B interactions, thus ultimately serving as a tool for modeling antibody production, mechanisms of immunological dysfunction, or toxicology in variable human populations.

Our data serve as a proof of principle that the LN-MPS can effectively model human T-B cell interactions and antibody production, and open the door to further mechanistic experiments into these events. For example, we used the strongest reported combination of Tfh skew conditions (IL-12 + Activin A)^3^ to maximize Tfh differentiation and increase the likelihood of inducing naïve B cells differentiation into plasmablasts / plasma cells and antibody secretion. Importantly, multiple cytokines (Activin A, IL-12, IL-6 and IL-23) have each separately been shown to induce the skewing of activated, naïve CD4+ T cells into pre-Tfh cells in humans.^3,71^ This diversity of cytokines clearly implicates redundancy in the system and further suggests that different cytokines may act at different times or locations within the lymph node. Furthermore, different combinations of cytokines may induce functionally different Tfh leading to different B cell differentiation outcomes (class-switching, hypermutation, etc.). It has been difficult to test such predictions in a human 3D lymphoid microenvironment previously. Similarly, while here we employed a maximal T cell stimulation (soluble anti-TCR + anti-CD28 for three days) and B cell stimulation (soluble anti-IgM + R848 for three days) for skewing, we anticipate that altering the duration or strength of these initial stimuli are likely to have major impacts on the subsequent B cell differentiation and antibody production. One of the greatest strengths of the LN-MPS model is that it provides an experimentally tractable system to directly investigate the impact of both the temporal or spatial modulation of these key signals on T cells and B cells. Future development of technologies that provide regulation of both the location and duration of T and B cell activation as well as the local cytokine milieu in the LN-MPS will enable a more complete and nuanced understanding of the underlying immunology that governs these key human antibody responses.

## METHODS

### Immunostaining and imaging of tonsil slices

Male and female human tonsils were provided by the UVA Biorepository and Tissue Research Facility as de-identified surgical discard tissue. The tonsil tissue was cleaned to remove portions damaged by surgery, and a 3 mm biopsy punch was used to obtain smaller pieces (Sigma Aldrich). 300-µm-thick tissue slices were collected and labelled for live immunofluorescence imaging according to procedures previously developed for murine lymph node tissues, with minor modifications as follows.^72,73^ Briefly, the small tonsil pieces were embedded in 6% w/v low melting point NuSieve GTG agarose (Lonza) in 1× PBS without calcium or magnesium (Gibco) and punched out with a 10-mm biopsy punch. The agarose was sterilized by autoclaving prior to embedding tissue for sterile slicing. The embedded tissue was sliced to 300 µm thickness using a Leica VT1000S vibratome (Leica) set to a frequency of 6 (60 Hz), amplitude of 5 (1 mm), and speed of 3.9 (∼0.153 mm/s). Immediately after slicing, the tissue slices were placed in sterile “RMPI complete media” composed of RPMI (Lonza, Maryland, USA) supplemented with 10% FBS (Corning, New York, USA), 1X L-glutamine (Gibco Life Technologies, Maryland, USA), 50 U/mL Pen/Strep (Gibco Life Technologies, Maryland, USA), 50 μM beta-mercaptoethanol (Gibco Life Technologies, Maryland, USA), 1 mM sodium pyruvate (Hyclone, Utah, USA), 1X non-essential amino acids (Hyclone, Utah, USA), and 20 mM HEPES (VWR, Pennsylvania, USA) 1X Normocin (InvivoGen, California, USA) and incubated in a cell culture incubator for at least 1 hour before staining. Figure a(i) tonsil came from unknown donor, and a(ii) tonsil came from a 3-year-old male African American/Non-Hispanic donor.

Following incubation, the tonsil slices were transferred to a Parafilm-covered surface, and a stainless-steel washer was placed on top. The tonsil tissue slices were treated with 25 µg/mL of anti-mouse CD16/32 in media for 30 minutes in a cell culture incubator. After blocking, 40 µg/mL fluorescently labeled antibody cocktail prepared in media was added to the slices, and they were incubated for an additional hour in the cell culture incubator. Finally, the stained tonsil slices were washed by immersion in sterile media for at least 30 min in a cell culture incubator before imaging. Tonsil slice imaging was performed on an upright Nikon A1Rsi confocal microscope, using 400, 487, 561 and 638 nm lasers with 450/50, 525/50, 600/50 and 685/70 nm GaAsP detectors, respectively. Images were collected with 4x and 40x/0.45NA Plan Apo NIR WD objectives. Image analysis was completed using ImageJ software 1.48v.

### Cell sourcing and culture media

Naïve, human CD4+ T cells and naïve B cells were purified from TRIMA collars, a derivative product of platelet apheresis. Donors were healthy and de-identified, with age and gender information reported (StemCell Technologies, Crimson Core, Brigham and Women’s Hospital, Boston, MA and INOVA Laboratories, Sterling, VA). Donor codes shown throughout the paper are “D” (for donor), “Age,” “Sex” (M/F, when reported). For example, a 77-year-old, female donor would be assigned the code “D77F.” Cells were purified using negative isolation. Briefly, whole blood from TRIMA collars was collected and diluted to a total volume of 30 mL using a 1X PBS 2% heat inactivated FBS solution. Diluted whole blood was evenly split and total T and B cell populations were enriched according to manufacturer protocol using RosetteSep Human T Cell Enrichment Cocktail (StemCell Technologies, Cat#15061) and RosetteSep Human B Cell Enrichment Cocktail (StemCell Technologies, Cat#15024), respectively. Following incubation, blood was further diluted to 35 mL total volume as previously described, overlayed over 15 mL of Ficoll-Paque Premium density gradient media, density 1.078 g/mL, (Cytiva, USA, Cat#17544202) and centrifuged at 1200G for 20 minutes with brake off. Following centrifugation, the top 25 mL of the plasma layer was discarded. The remaining plasma layer, interphase layer, and density gradient media layer was collected, leaving coagulated pellet undisturbed. Collected solution s were washed twice and resuspended in 1X PBS, 2% heat inactivated FBS, and 1 mM EDTA in preparation for naïve cell isolation. Naïve CD4+ T and CD19+ B cell populations were isolated using EasySep Human Naïve CD4+ T cell Isolation Kit (StemCell, Cat#19555) and EasySep Human Naïve B Cell Isolation Kit (StemCell, Cat#17254) following manufacturer protocols. Analysis of isolation efficiency by flow cytometry confirmed on average >90% pure naïve CD4+ T cells and >90% pure naïve CD19+ B cells were isolated.

Cells were cultured in serum-free AIM-V media containing phenol red pH indicator, L-glutamine, 50 μg/mL streptomycin sulfate and 10 μg/mL gentamicin sulfate (Gibco, P/N 12055083). In 2D culture, T and B lymphocytes were cultured at 1-3 x 10^6^ cells/mL. For maintaining viable naïve CD4+ T cells, IL-7 was added at a concentration of 4 ng/mL. To activate naïve CD4+ T cells in 2D and 3D culture, ImmunoCult™ Human CD3/CD28 T Cell Activator (StemCell Technologies) was added to the media at a concentration of 25 μL/mL media. For Tfh skewing, naïve CD4+T cells were activated with ImmunoCult™ and cultured with IL-12 (5 ng/mL) and activin A (100 ng/mL) (R&D Systems). To activate naïve B cells in monoculture, a cocktail of human α-IgG/IgM (10 µg/mL) (Affinipure Goat Antihuman IgG+IgM (H+L), Jackson ImmunoResearch), R848 (2 µg/mL) (Invivogen), and CD40 ligand (100 µg/mL) was added to the culture media. When B cells were activated in co-culture with T cells, CD40 ligand was not added to the cocktail. Staphylococcus enterotoxin B (SEB) (Toxin Technology) was added at a concentration of 1 µg/mL.

### Hydrogel preparation

Collagen/fibrinogen hydrogel was prepared by diluting 5 mg/mL rat tail I collagen (Ibidi) and 2 mg/mL fibrinogen (Sigma Aldrich) in 1x PBS. The pH was adjusted to 7.4 while on ice, according to the Ibidi protocol for preparation of collagen gels, to a final concentration of 1.5 mg/mL collagen and 1 mg/mL fibrinogen. For experiments involving T or B lymphocytes in isolation on chip, cells were resuspended in hydrogel at 1.0x10^7^ cells/mL. For experiments with T/B co-cultures, cells were resuspended in hydrogel at a density of 2.5x10^7^ cells/mL.

### Microfluidic chip design and fabrication

The microfluidic housing used for the LN MPS was fabricated from a thin layer of polydimethylsiloxane (PDMS) patterned by standard soft lithography and irreversibly bonded to a 50 x 75 mm glass slide.^51^ The PDMS microchamber was comprised of five parallel channels separated by arrays of hexagonal micropillars (100 µm in vertex-to-vertex diameter, with 50 µm spacing between pillars). The outer channels each had 8-mm diameter inlets and outlets that served as media reservoirs, while the inner gel channels had 0.75-mm inlets and outlets. Excluding the converging, angled channel regions, the media lanes were 2.05 x 0.13 x 3.5 mm^3^, and the gel lanes were 0.3 x 0.13 x 3.5 mm^3^. A schematic is included in the supplemental information (Figure S4).

### Viability staining on chip

Viability staining was performed by first removing bulk media from the media reservoirs and rinsing each side of the chip with 150 μL of 1X PBS. Bulk rinsate was removed from the wells and 150 μL of 10 μM Calcein AM and 1 μM DAPI solution in 1X PBS was added to each side of the chip (Fisher Scientific). Chips were placed in a cell culture incubator for 1 hr. The media reservoirs were then rinsed twice with excess (500 μL) PBS, once every thirty minutes, for 1 hr., and then imaged immediately.

### Chemotaxis measurement on chip

For assessing chemotactic activity of naïve CD4+ T cells on chip, a solution of 0.1 μM CCL21 (recombinant human, Peprotech, NJ, USA, Cat# 300-35A) in AIM-V media was added to the left media reservoir. The chip was incubated in a cell culture incubator for 30 min for assessing live-cell motility or 1 hr for assessing total cell displacement. Cells were imaged using brightfield microscopy (no labeling) for live-cell imaging, and cells were labeled with 10 µM Calcein AM prior to imaging for total cell displacement. To perform live-cell motility imaging, chips were imaged on a stage-top incubator set to 37°C at 30 sec intervals for 5 min. For calculating mean cell velocity, individual cells were tracked using CellTracker ver.1.1 with semi-automated tracking; matching modality was set to histogram matching, maximal cell displacement was set to 20, and cell diameter was set to 40.^74^

### Immunofluorescence staining on chip

For immunofluorescence staining on chip, the bulk culture media was removed from the media reservoirs, and chips were rinsed with 150 μL of 1X PBS. Bulk rinsate was removed from the wells and 100 μL of the appropriate antibody cocktail prepared in Hanks Buffered Saline Solution (ThermoFisher, Cat# 14025092) was placed in the top media reservoirs. Antibody information is available in Table S1. Antibody cocktail was allowed to flow to the bottom media reservoirs via gravity-driven flow. Cells were stained for 1 hr. while incubating at 37 °C, and then the media reservoirs were rinsed first with HBSS for 20 min. and then twice with AIM-V media for 20 min. prior to imaging.

### Microscopy of the LN MPS

All microscopy performed within the LN MPS was done using either a ZEISS AxioZoom.V16 or ZEISS AxioObserver Inverted. The AxioZoom.V16 was fitted with a HXP 200C metal halide illumination source, PlanNewFluor Z 1X objective (0.25 NA, FWD 56 mm), and an Axiocam 506 mono camera. Fluorescence imaging used Zeiss Filter Sets 38 HE (Ex: 470/40, Em: 525/50); 43 (Ex: 550/25, Em: 605/70); 49 HE (Ex: 365, Em: 445/50); 64 HE (Ex: 587/25, Em: 647/70); and brightfield images collected using transmitted light. The AxioObserver Inverted was fitted with a Colibri.7 LED light source, LD PN 20X objective (0.4 NA, FWD 7.9 mm), and ORCA-Flash4.0 LT + sCMOS camera (Hamamatsu). Transmitted light was used for collecting brightfield images and a ZEISS 112 HE LED penta-band filter was used for acquiring fluorescent images. Zen 3 Blue software was used for image collection. Image analysis was completed using ImageJ software 1.48v.

### Cytokine and antibody secretion detection

All cytokines and antibodies were detected by sandwich ELISA in a high binding 96-well plate (Fisher Healthcare, Cat# 07-200-37). IgM was detected using a Human IgM ELISA Antibody Pair Kit (StemCell technologies, Cat#01995), and IL-21 was detected using an ELISA Flex: Human IL-21 (HRP) kit (Mabtech, Ohio, USA, Cat#3540-1H-6) following manufacturer protocols. IFN-γ was detected using an in-house optimized ELISA assay. Briefly, unconjugated mouse monoclonal anti-human IFN-γ (clone NIB42) was used as the capture antibody, recombinant human IFN-γ (peprotech, Cat# 300-02) was used as the standard, and biotinylated mouse monoclonal anti-human IFN-γ (clone 4S.B3) was used as the detection antibody. 150 μL/well ELISA wash buffer (1% BSA, 0.05% TWEEN-20, 1X PBS) was used to rinse plates in-between all steps. 150 μL/well blocking buffer (1% BSA, 1X PBS) was used to block plates, and all other reagents were added at 50 μL/well. Well plates were incubated overnight at 4 °C using 1 μg/mL capture antibody in 1X PBS. Following rinsing (3X), plates were blocked with ELISA block buffer for 1 hour at room temperature. Plates were rinsed again (3X) and samples and standards were added to the plate for a 1.5 hr. room-temp incubation. Detection antibody was prepared at 0.5 μg/mL in ELISA block buffer, added after rinsing (3X), and incubated for 1.5 hrs. at room temp. Avidin-HRP (biolegend, Cat# 405103) was diluted in ELISA block buffer at a 1:500 ratio, added to the plate following rinsing (3X), and incubated for 30 min. at room temperature, in the dark. The plate was then rinsed again (5X) and TMB substrate (Fisher Healthcare, Cat# BDB555214) was added to the plate, and incubated in the dark until a color gradient across the standard wells was observed, no more than 10 min. The TMB-HRP reaction was stopped using 1M H_2_SO_4_ and absorbance readings taken at 450 nm on a plate reader (BMG Labtech Clariostar).

### Cell recovery for flow cytometric analysis

To recover cells, collagenase D (1 mg/mL in PBS) (Sigma Aldrich) was added to the media lanes and incubated at 37 °C for fifteen minutes. Following digestion, the elastic PDMS housing was massaged with a pipette tip to break up the internal 3D cell culture. Finally, an ice-cold solution 2% FBS in 1X PBS was used to rinse across the media lanes. To recover enough cells for analysis, cells were pooled from up to twenty chips from a single donor, as indicated in the figure captions.

### Flow cytometry

Cells were incubated with the fixable viability dye efluor 780 (eBioscience, ThermoFisher Scientific) according to manufacturer’s instructions to determine the overall viability of lymphocytes in culture. To assess the differentiation state of T and B lymphocytes, cells were suspended in FACS staining buffer (PBS + 2% FBS—Heat Inactivated + 0.1% Sodium Azide) and first stained with biotin-conjugated rat anti-human CXCR5 antibody (Clone: RF8B2, BD Biosciences). After staining with anti-CXCR5 antibody, the cells were stained with the following fluorescently conjugated reagents: Brilliant Violet 421 conjugated streptavidin, anti-human CD69 (FITC), anti-human CD4 (PE-Cy7), anti-human CD38 (PE), anti-human CD19 (Alexa Fluor 647), anti-human PD-1 (Brilliant Violet 711), and anti-human CD20 (Brilliant Violet 510). All the previously listed antibodies were purchased from Biolegend.

For intracellular staining, cells were fixed and permeabilized after staining for surface antigens using the Foxp3 intracellular staining kit (eBioscience, ThermoFisher Scientific) according to manufacturer’s instructions. Permeabilized cells were then stained with either Alexa Fluor 647 conjugated mouse anti-human BCL6 (clone: K112-91, BD Biosciences) or mouse IgG1,κ isotype control (Biolegend).

Stained cells were acquired using the Attune NXT flow cytometer (ThermoFisher Scientific) and flow cytometry data were analyzed using FlowJo software (BD Biosciences) and GraphPad Prism.

Pre-Tfh cells were defined as CD4^+^CXCR5^hi^PD-1^hi^ (surface expression) or CD4^+^CXCR5^hi^BCL6^hi^ cells (intracellular staining), and plasmablasts were defined as having a surface expression phenotype of CD19^+^CD38^hi^CD20^lo^ cells.

### Statistical analysis

Graphs, line-fitting, and statistical analysis was prepared using GraphPad Prism 8.4.2.

Specific statistical tests used for each figure are available in the figure captions.

## DATA AVAILABILITY STATEMENT

The data that support the findings of this study are available from the corresponding author upon reasonable request.

## AUTHOR CONTRIBUTIONS

JMZ, DR, AA, TO, JOC, CJL, JMM, and RRP led the experimental design and interpreted the findings. JMZ, DR, AA, SK, TO, PA, and RRP drafted the manuscript. JMZ designed and produced the microfluidic device and related experiments for 3D culture, manipulation, and characterization of lymphocytes. AA designed lymphocyte culture protocols, isolated and performed culture experiments with lymphocytes in 2D, and designed and performed all flow cytometry characterization experiments. DR designed and conducted the co-culture experiments on chip, with technical assistance from SK. DR and TO contributed to experimental set-up and data collection, and performed ELISAs. PA performed tonsil slicing and its immunofluorescence staining and imaging. JMM provided tissue engineering and tissue analysis expertise and edited the manuscript. CJL provided immunology expertise regarding lymphocyte isolation, activation, and co-culture. TJB provided immunology expertise regarding the various strategies for activating lymphocytes. The manuscript was critically revised by all authors with the exception of TJB, who sadly passed away prior to completion of this study.

## ACKNOWLEDGEMENTS

This work was supported by the National Institute of Biomedical Imaging and Bioengineering (NIBIB), with co-funding from the National Center for Advancing Translational Sciences (NCATS) at the National Institutes of Health (NIH), under Award Number U01EB029127. JMZ was supported in part by the National Science Foundation Graduate Research Fellowship Program (NSF GRFP). TO was supported in part by a Summer Research Award through the Global Infectious Diseases Institute at the University of Virginia. PA was supported by the National Institute of Allergy and Infectious Diseases under Award Number R01AI131723. JOC was supported in part by a training grant from the National Institute of General Medical Sciences of the National Institutes of Health under Award Number T32GM136615. JHH was supported by the Institute for Critical Technology and Applied Science (ICTAS) at Virginia Tech. The content of this paper is solely the responsibility of the authors and does not necessarily represent the official views of the National Institutes of Health.

## SUPPLEMENTARY DATA

Please see the SI document for supporting formation which includes Figure S1. Schematic of the master mold used in patterning the LN-MPS, Figure S2. Timeline of naïve T and B cell viability in 2D monoculture, with and without cytokine supplementation, with accompanying gating strategy, Figure S3. Naïve B cell activation on chip, Figure S4. Gating strategies pre-Tfh characterization, Figure S5. ITS and ITS-X supplement improves viability of T-B co-cultures, with accompanying gating strategy, Figure S6. Time course of IgM secretion as a function of T-B proximity, Figure S7. Brightfield imaging time course displaying gradual increase to cell density and cluster formation, Figure S8. Impact of T cell phenotype on B cell plasmablast differentiation and IgM secretion, Table S1. List of antibodies used for fluorescent microscopy, and Video S1. Looped video of T and B lymphocytes moving through the MPS.

