## Supplemental Files for "Initiation of primary T cell—B cell interactions and early antibody responses in an organized microphysiological model of the human lymph node"

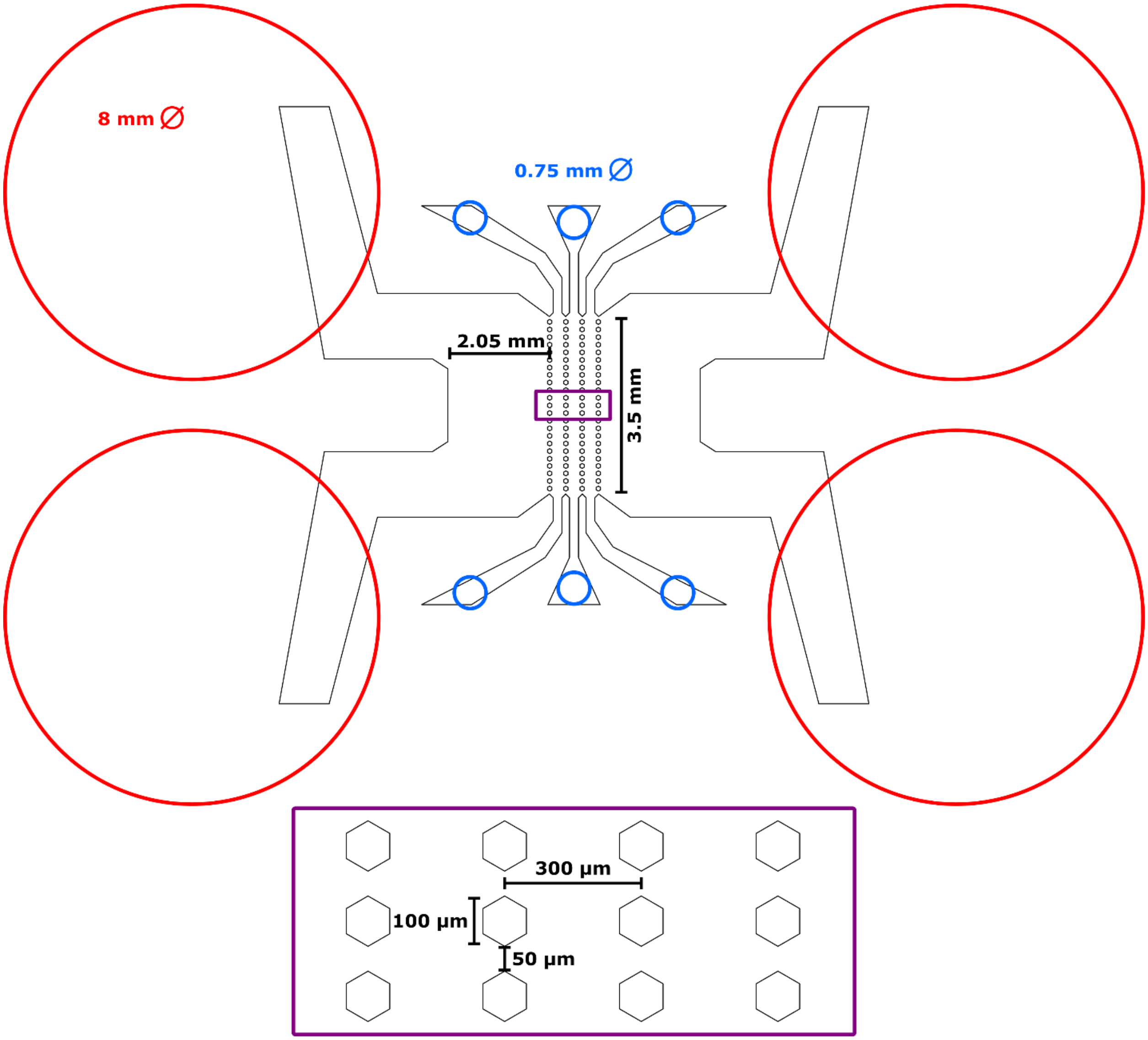

**Figure S1. Schematic of the master mold used in patterning the LN-MPS.** A 2D schematic (drawn to scale) showing the dimensions of the master mold used for patterning the T-B border chip in PDMS. Large red circles indicate where tissue biopsy punches were used to cut out reservoirs for cell culture media. Smaller blue circles indicate where tissue biopsy punches were used to cut out inlets/outlets for loading collagen-fibrinogen hydrogel, either with or without cells.

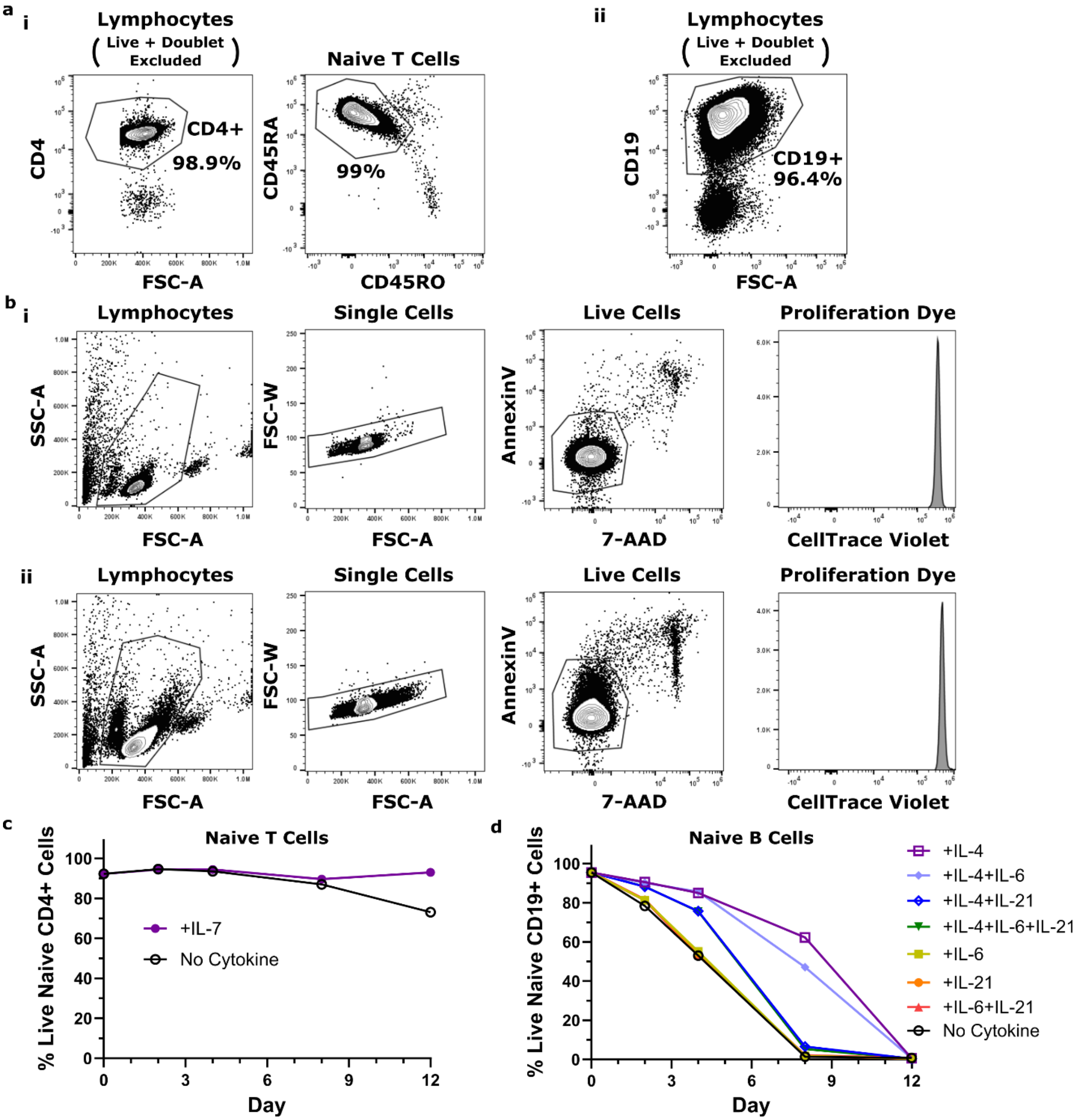

**Figure S2. Purity and viability of naïve human T and B cells in 2D monoculture, with and without cytokine supplementation.** Purified populations were cultured in AIM-V media without ITS supplementation at 1 x $10^6$ cells/mL. (a) Gating strategies for determining purity of (i) naïve CD4+ T cell and (ii) naïve B cell isolates. (b) Gating strategies for determining impact of cytokine supplementation on viability of purified naïve (i) CD4+ T cells and (ii) B cells. (c) Viability of purified naïve CD4+ T cells with and without IL-7 supplementation. (d) Viability of purified CD19+ naïve B cells with and without cytokine supplementation. IL-7 improved naïve T cell viability, and IL-4 improved naïve B cell viability in classic 2D monoculture.

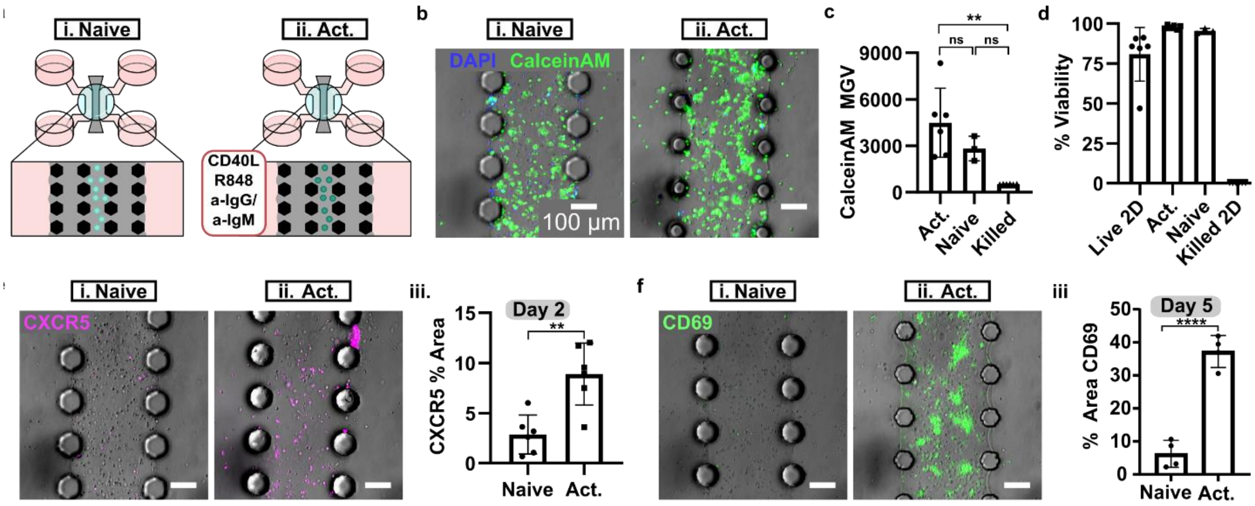

**Figure S3. Naïve B cell activation on chip.** Purified naïve B cells were loaded into the LN-MPS and cultured with or without an activation cocktail composed of CD40L, R848, and anti-IgG/IgM. (a) Schematic of naïve (i) and activated (ii) culture conditions on-chip for purified naïve CD19+ B cells. (b) Representative images of naïve (i) and activated (ii) B cells on chip after 48 hours of culture. Stained with viability dyes; Calcein AM, green, live; DAPI, blue, dead. (c) Quantification of Calcein AM mean grey value across the whole image. Ordinary one-way ANOVA with Tukey's multiple comparisons, $^{**}p<0.005$. (d) Quantification of percent viability on chip and in off-chip (2D) controls. (e) Representative images of naïve (i) and activated (ii) B cells on chip after immunofluorescence staining with AlexaFluor647-anti-CXCR5 (magenta). (iii) Quantification of percent positive CXCR5 area across the whole image. Unpaired T test, $^{**}p<0.005$. (f) Representative images of naïve (i) and activated (ii) B cells on chip stained with FITC-anti-CD69 (green). (iii) Quantification of percent positive CD69 area across the whole image. Unpaired T test, $^{****}p<0.00005$. Each dot is a chip. Donor information: D76M (b-e), D24F (f).

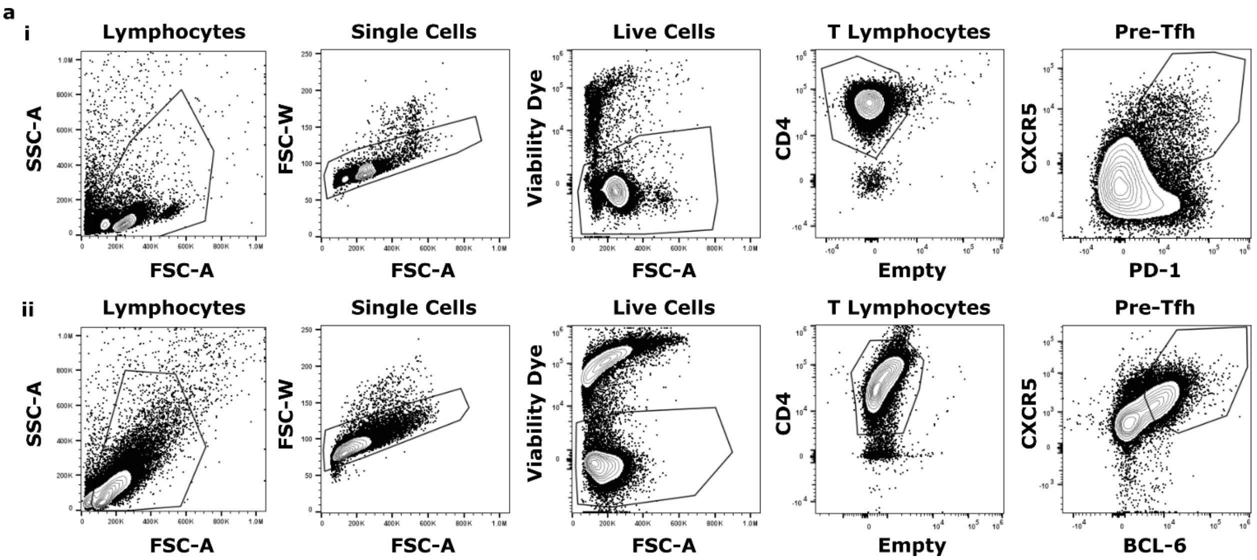

**Figure S4. Gating strategies for pre-Tfh characterization shown in Figure 3.** Purified naïve CD4+ T cells were cultured in the LN-MPS with or without a skewing cocktail (IL-7, IL-12, αCD3/CD28, and Activin A). Cells recovered from the chips were analyzed by flow cytometry. (a) Gating strategy for measuring degree of pre-Tfh development, defined as (i) CXCR5+ and PD-1+ cells, or (ii) CXCR5+ and BCL-6+ cells. The viability dye used was eBioscience™ Fixable Viability Dye eFluor™ 780.

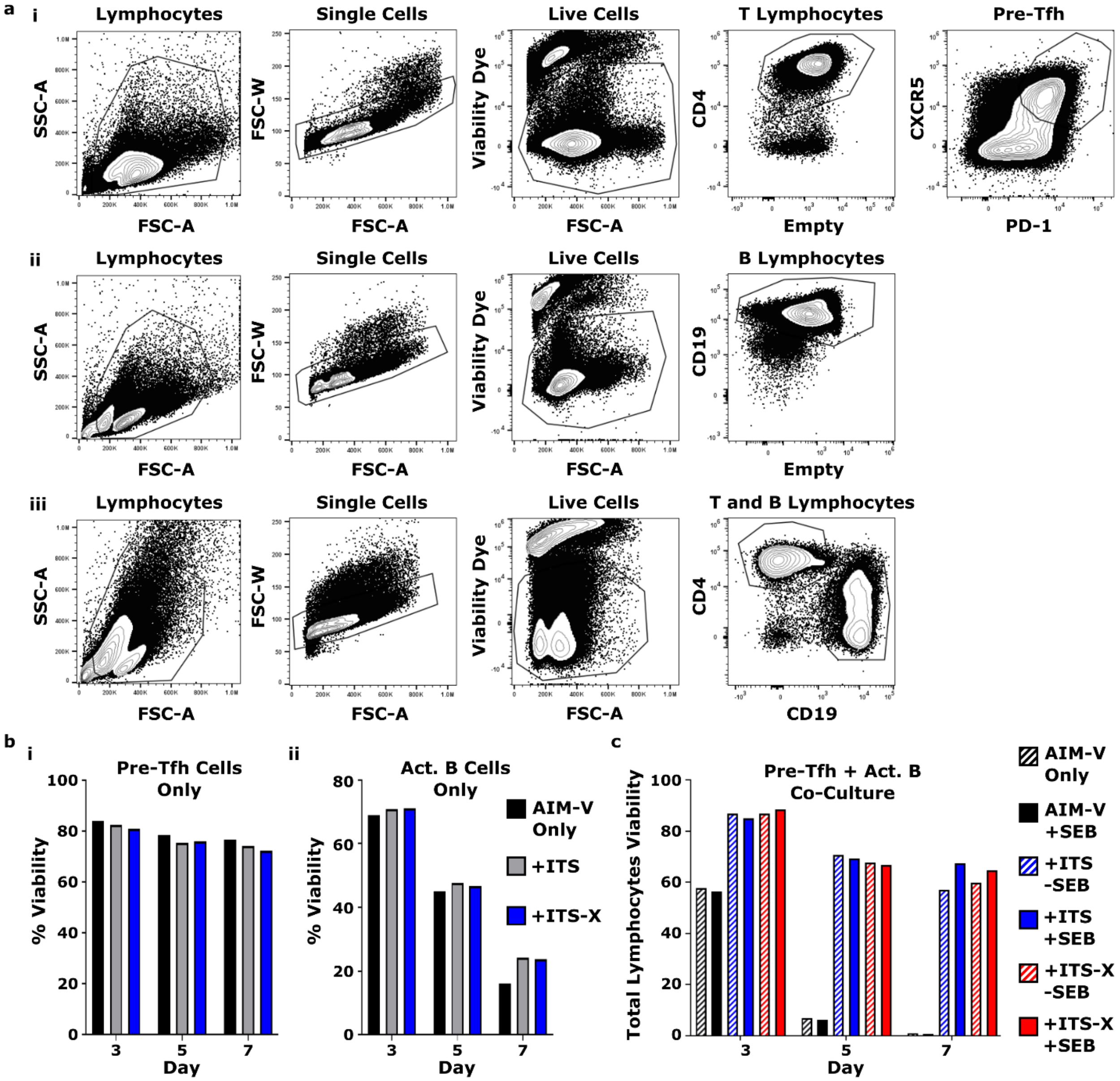

**Figure S5. ITS and ITS-X supplementation improves viability of T-B co-cultures.** T and B cells were cultured either individually or together in the presence or absence of ITS/ITS-X. (a) Gating strategy for measuring viability of pre-Tfh cells (i) and activated B cells (ii) in monoculture, and pre-Tfh cells and activated B cells in co-culture (iii). (b-i) Plot showing viability of naïve T cells after 3 – 7 days of culture in a pre-Tfh skewing cocktail (IL-7, IL-12, Activin A, anti-CD3/CD28) and (b-ii) plot showing naïve B cell viability after 3 – 7 days of culture in an activating B cocktail (CD40L, R848, anti-IgG/IgM) (ii). (c) Plot showing total lymphocyte viability over time, for pre-Tfh-skewed T cells and activated B cells after 2 – 7 days of co-culture. Cells were skewed individually for three days prior to co-culture. All results are from 2D cultures. Viability determined by flow cytometry using Fixable Viability Dye eFluor 780. One donor, D56F.

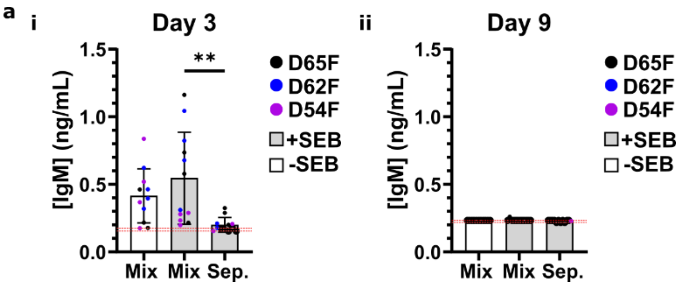

**Figure S6. Additional time points for IgM secretion as a function of T-B proximity and SEB, accompanying Figure 4.** IgM secretion was not SEB dependent on day 3, and IgM stopped being secreted by day 9. (a) Quantification of IgM secretion for days 3 (i) and 9 (ii) as measured by ELISA. Each dot is a single chip replicate and error bars show the standard deviation of the response across all three donors for a given condition. Statistics determined using an ordinary on-way ANOVA with Sidak's multiple comparisons test with single pooled variance. **$p<0.003$. Dashed red line is the limit of detection.

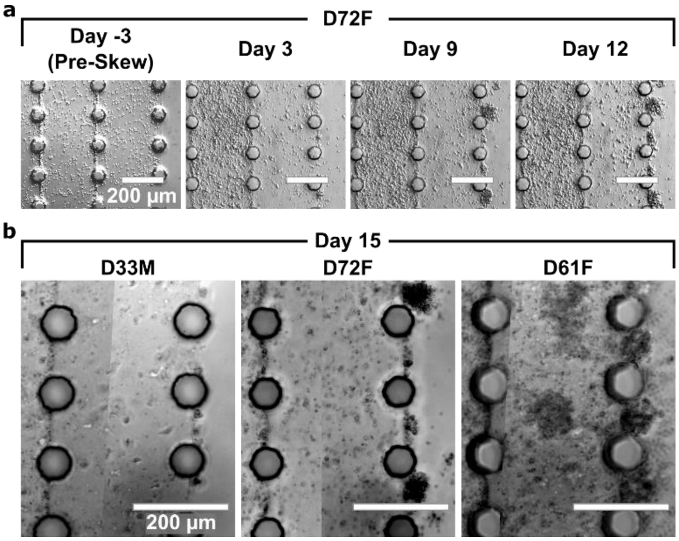

**Figure S7. Brightfield imaging time course displaying gradual increase in cell density and cluster formation over time.** Naïve T and B cells were skewed in co-culture in the LN-MPS and respond to SEB with varying degrees of increase to cell density and cluster formation. (a) Representative brightfield images of the middle and right-most gel lanes, adjacent to a media lane, showing increase in cell density in the gel lanes and clusters forming at the gel-media interface over time. (b) Representative brightfield images of right-most gel lane next to media lane, from day 15, for three donors, showing variable responses to SEB stimulation.

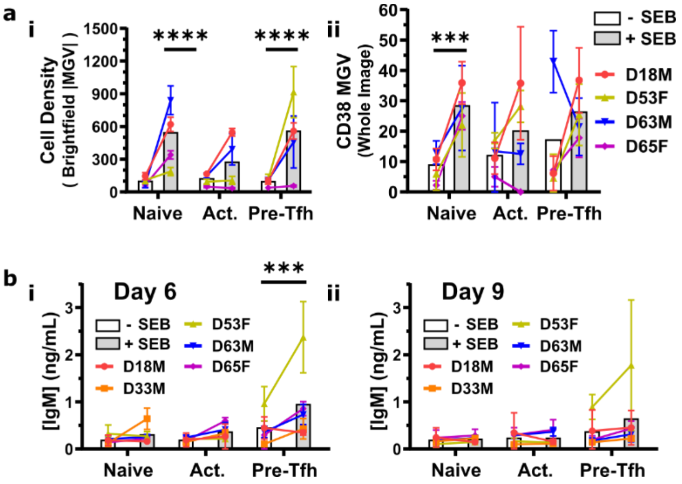

**Figure S8. Impact of T cell phenotype on B cell plasmablast differentiation and IgM secretion.** SEB-dependent IgM secretion is dependent on the T cell phenotype but CD38+ plasmablast formation is not. (a) Quantification of increased cell density (i) at end-point analysis in response to SEB and T cell phenotype using the absolute value difference between the MGV of the empty media lanes and cell-populated gel lanes. (ii) Quantification of percent positive CD38 area across the whole image in response to SEB and T cell phenotype. (b) Quantification of IgM secretion in response to SEB and T cell phenotype on day six (i) and day nine (ii). Statistics for all panels determined using ordinary two-way ANOVA with Šidák's multiple comparisons test and single pooled variance, *** p=0.0005, **** p<0.00001. Each dot is the mean response for a given donor measured from 2-4 individual chips, error bars show the standard deviation for a given donor, and background bars show the combined mean response.

**Table 1. List of antibodies used**

| Host Species | Conjugate | Immunogen | Clone | Working Conc. | Vendor | Cat# |
|---|---|---|---|---|---|---|
| ***For Microscopy*** | | | | | | |
| Rat | FITC | Podoplanin | NC-08 | 40 µg/mL | Biolegend | 337026 |
| Mouse | PE | CD19 | SJ25C1 | 40 µg/mL | Biolegend | 363004 |
| Mouse | AF647 | CD38 | HIT3a | 40 µg/mL | Biolegend | 300322 |
| Mouse | FITC | CD69 | FN50 | 1 µg/mL | Biolegend | 310904 |
| Rat | AF647 | CXCR5 | RF8B2 | 2 µg/mL | BD Pharmigen | 558113 |
| Mouse | BV421 | CD4 | RPA-T4 | 1 µg/mL | Biolegend | 300532 |
| Mouse | DL488 | CD138 | MI15 | 2 µg/mL | Biolegend | 356502 |
| Mouse | AF546 | CD38 | AT-1 | 2 µg/mL | Santa Cruz Biotechnology | sc-7325 AF546 |
| Mouse | AF647 | CD19 | F-3 | 2 µg/mL | Santa Cruz Biotechnology | sc-373897 AF647 |
| ***For Flow Cytometry*** | | | | | | |
| Rat | Biotin | CXCR5 | RF8B2 | 1:20 | BD Biosciences | 552118 |
| Mouse | AF647 | Bcl6 | K112-91 | 1:20 | BD Biosciences | 561525 |
| Mouse | PE | CD38 | S17015A | 1:20 | Biolegend | 397104 |
| Mouse | FITC | CD69 | FN50 | 1:100 | Biolegend | 310904 |
| Mouse | BV605 | CD69 | FN50 | 1:100 | Biolegend | 310938 |
| Mouse | BV711 | CD20 | 2H7 | 1:20 | Biolegend | 302342 |
| Mouse | BV510 | CD20 | 2H7 | 1:20 | Biolegend | 302340 |
| Mouse | APC | CD20 | 2H7 | 1:20 | Biolegend | 980206 |
| Mouse | PE-Cy7 | CD4 | SK3 | 1:100 | Biolegend | 344612 |
| Mouse | BV711 | PD-1 | NAT105 | 1:20 | Biolegend | 367428 |
| Mouse | BB515 | PD-1 | EH12.1 | 1:20 | BD Biosciences | 564494 |
| Mouse | BV421 | CD19 | SJ25C1 | 1:100 | Biolegend | 363018 |
| Mouse | AF647 | CD19 | SJ25C1 | 1:100 | Biolegend | 363040 |

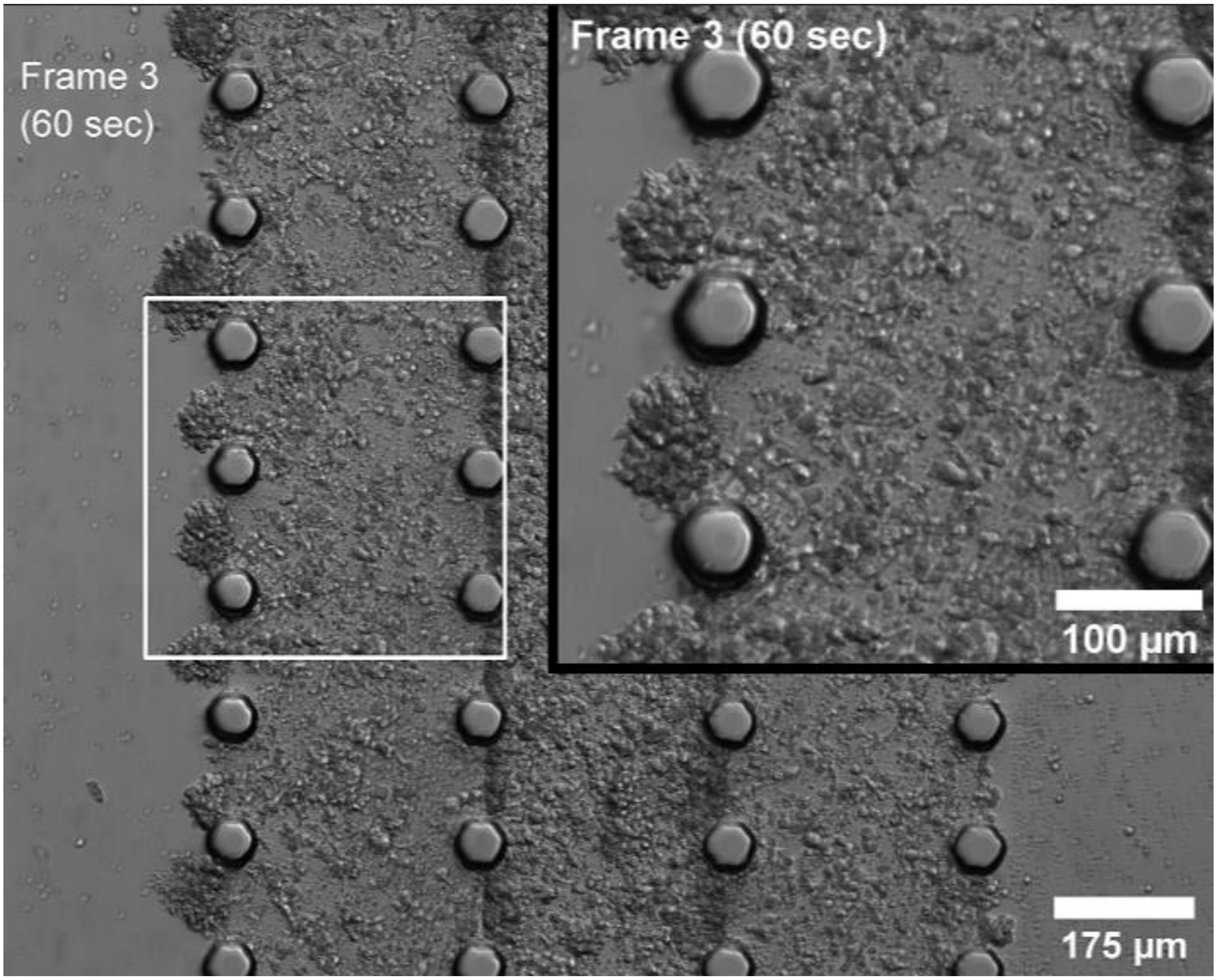

**Video 1. Looped video of T and B lymphocytes moving through the MPS.** Pre-Tfh and activated B cells were seeded on chip at a 1:5 ratio and total cell density of 2.5 x $10^7$ cells/mL. Culture media containing SEB was refreshed every three days and the LN-MPS was imaged on day 9. Images were taken every 30 seconds for five minutes and compiled into a video, single video frame shown above.
